# CRISPR-Cas9 Screening Reveals Microproteins Regulating Adipocyte Proliferation and Lipid Metabolism

**DOI:** 10.1101/2025.03.21.644636

**Authors:** Victor J. Pai, Hazel Shan, Cynthia Donaldson, Joan Vaughan, Carolyn O’Connor, Michelle Liem, Antonio Pinto, Jolene Diedrich, Alan Saghatelian

## Abstract

Small open reading frames (smORFs) encode microproteins that play crucial roles in various biological processes, yet their functions in adipocyte biology remain largely unexplored. In a previous study, we identified thousands of smORFs in white and brown adipocytes derived from the stromal vascular fraction (SVF) of mice using ribosome profiling (Ribo-Seq). Here, we expand on this work by identifying additional smORFs related to adipocytes using the *in vitro* 3T3-L1 preadipocyte model. To systematically investigate the functional relevance of these smORFs, we designed a custom CRISPR/Cas9 guide RNA (sgRNA) library and screened for smORFs influencing adipocyte proliferation and differentiation. Through a dropout screen and fluorescence-assisted cell sorting (FACS) of lipid droplets, we identified dozens of smORFs that regulate either cell proliferation or lipid accumulation. Among these, we validated a novel microprotein as a key regulator of adipocyte differentiation. These findings highlight the potential of CRISPR/Cas9-based screening to uncover functional smORFs and provide a framework for further exploration of microproteins in adipocyte biology and metabolic regulation.

**Significance:** Obesity and its associated metabolic disorders pose significant public health challenges, yet the molecular mechanisms regulating adipocyte function remain incompletely understood. Small open reading frames (smORFs) and their encoded microproteins represent an emerging class of regulatory elements with potential roles in metabolism. Here, we leveraged CRISPR/Cas9 screening to functionally characterize smORFs in adipocytes, identifying novel regulators of cell proliferation and lipid metabolism. Our findings demonstrate that conservation is not a prerequisite for smORF function, as we validated a mouse-specific microprotein that modulates adipocyte differentiation. This work establishes a robust pipeline for unbiased smORF discovery and highlights the potential for species-specific microproteins to regulate adipose biology. Future studies in human adipocytes may uncover additional microproteins with therapeutic relevance for obesity and metabolic disease.

## Introduction

Algorithmic analyses of genome sequences revealed hundreds of thousands to millions of small open reading frames (smORFs) in eukaryotes (1). However, these smORFs were intentionally excluded from databases because they were thought to be derived by chance from the random assembly of codons in genomes, and most of these smORFs are not conserved—raising doubts about their evolutionary significance (2–7). Also, smORFs often appeared in the untranslated regions (5′- and 3′-UTRs) of mRNAs already containing a longer translated ORF, challenging the long-held belief that each mRNA gives rise to a single protein (2, 3, 8). In the absence of methods to confirm the translation of smORFs, the contribution of smORF translation to the proteome remained enigmatic.

The advent of ribosome profiling (Ribo-Seq) revolutionized our understanding of translation. Ribo-Seq uses ribosome footprinting on RNA to identify regions of active RNA translation (9, 10). This technique provided an unbiased method to establish translated smORFs and revealed thousands of previously unannotated protein-coding smORFs in many genomes (11, 12). Many of these smORFs originated from RNAs or RNA regions previously considered non-coding, including non-coding RNAs and the 5’ and 3’-UTRs of mRNAs. These discoveries challenged the definition of “non-coding” and the conventional view that a single mRNA transcript produces only one protein (13). Subsequent proteomics studies, using custom databases designed to identify non-canonical translation products (i.e., proteogenomics), confirmed the translation of hundreds of smORF into stable peptides and small proteins. These were subsequently termed microproteins or micropeptides (8, 13). Together, these methods expanded the size of proteomes by 10-30% and revealed a rich source of unstudied proteins.

Many labs have begun uncovering functions for microproteins. For example, a family of single-pass transmembrane microproteins in muscle interacts with the transmembrane domain of sarcoplasmic/endoplasmic reticulum Ca^2+^-ATPase (SERCA) to regulate calcium flux and physiological traits such as muscle endurance in different muscle depots(14). A literature survey reveals diverse biological functions for microproteins in cells and tissues, including roles in DNA repair (15, 16), mitochondrial function (17–19), and cancer (20–22). Nevertheless, only a tiny fraction of all microproteins have been characterized to date, leaving the possibility for the continued discovery of many functional microproteins.

Of particular interest here is the role of microproteins in adipocyte biology. Obesity remains a significant public health challenge, associated with comorbidities such as type 2 diabetes and fatty liver disease (23, 24). Current FDA-approved treatments for obesity, including pharmacological interventions and bariatric surgery, primarily target reductions in overall energy intake (25–27). Peptide therapeutics, which can be considered to be microproteins, such GLP-1 receptor agonists like Semaglutide and Tirzepatide have shown exceptional efficacy in promoting weight loss (28), and provide support for the discovery of additional microproteins. Further investigation into adipocyte biology could reveal novel mechanisms for regulating adipose tissue and identifying new therapeutic strategies for obesity management.

Many functional microproteins were characterized because of the evolutionary conservation of their amino acid sequence (29, 30). Focusing solely on highly conserved microproteins could simplify the investigation of these genes in adipocyte biology but would overlook many potential candidates. We hypothesize that employing unbiased, experimental strategies to select smORFs for validation can reveal functional microproteins that might otherwise be dismissed. An example of this approach comes from a CRISPR screen combined with Perturb-Seq, which has successfully identified numerous microproteins, including non-conserved ones, that influence gene expression and cellular proliferation (31). However, while CRISPR screening enables large-scale phenotypic validation of smORF function, its observed effects could arise through multiple mechanisms. For instance, smORFs that overlap 5’ and 3’ UTRs of annotated genes can regulate the translation of the primary ORF on the same mRNA, as previously noted (32).

Additionally, CRISPR-induced disruptions in the 5’ or 3’ UTR regions could inadvertently impact the expression of the main annotated gene. Similarly, for non-coding RNAs, one possibility is that the observed phenotype—wholly or in part—results from knocking out the non-coding RNA rather than the microprotein itself (33). To confirm that the encoded microprotein is the functional genetic element, additional studies expressing the microprotein in trans must be conducted.

In a previous study, we used Ribo-Seq to identify thousands of smORFs in white and brown adipocytes derived from the stromal vascular fraction (SVF) of mice (SVF smORFs) (12). Here, we extend our Ribo-Seq data to identify additional smORFs in 3T3-L1 cells (3T3-L1 smORFs). The 3T3-L1 mouse adipocyte model is a widely used system for studying adipogenesis and fat cell biology, as it faithfully replicates the key differentiation steps from preadipocytes to mature adipocytes. This model provides a controlled and reproducible platform for exploring molecular mechanisms underlying adipocyte function, metabolism, and obesity-related disorders. Using CRISPR/Cas9 with a custom guide RNA (sgRNA) library, we investigated the activity of SVF and 3T3-L1 smORFs in 3T3-L1 adipocyte proliferation and differentiation. Our approach identified smORFs essential for adipocyte proliferation through a dropout screen and a fluorescence-assisted cell sorting (FACS) assay that stains lipid droplets to pinpoint smORFs modulating adipocyte differentiation (Fig. 1). In total, we discovered dozens of smORFs affecting either cell proliferation or lipid droplet formation. We validated a novel microprotein regulator of adipocyte differentiation. These findings lay a strong foundation for future CRISPR/Cas9-based screenings to interrogate smORF functions across diverse phenotypes and biological contexts.

**Figure 1.**
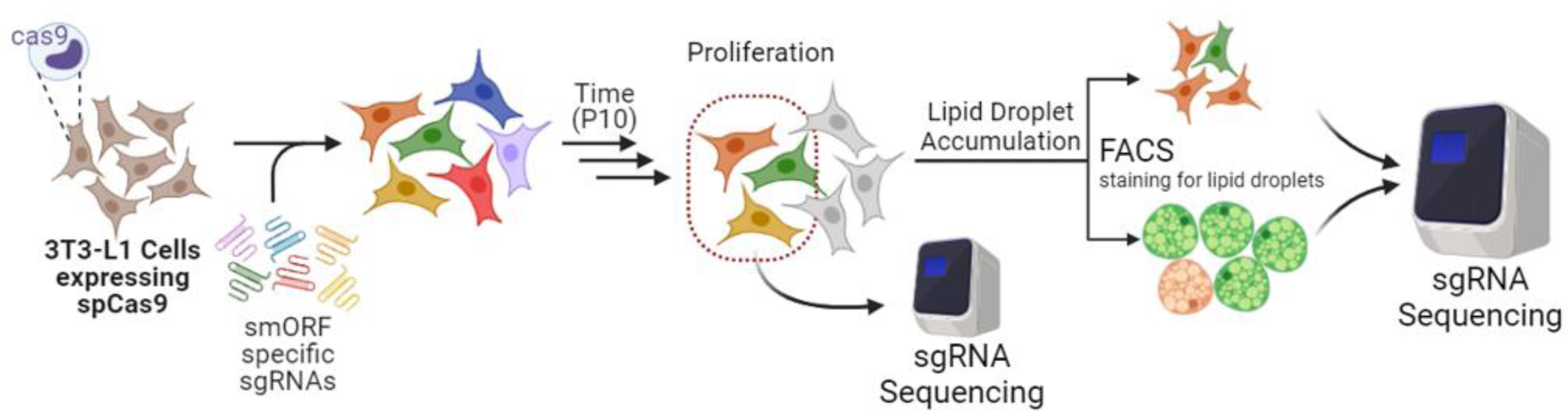
Experimental outline of CRISPR-Cas9 screen targeting adipocyte lipid droplet size in 3T3-L1 Cells. 3T3-L1 cells expressing spCas9 are transduced by a custom lentiviral sgRNA library targeting smORFs. After 10 Passages (30 days) cells are collected for sequencing or induced to become mature adipocytes by differentiation cocktail. Cells are stained for lipid droplets by BODIPY 493/503, and BODIPY^low^ and BODIPY^high^ cells are collected by FACS for sequencing.

## Results

### Ribo-Seq Elucidation of 3T3-L1 smORFs

To compile a list of candidate microproteins potentially involved in adipocyte function, we began with a previously published mouse adipocyte smORF dataset derived from ribosome profiling of smORFs in the stromal vascular fraction (SVF) (12). We only included smORFs consistently replicated across different ribosome profiling runs, narrowing the list to 944 SVF smORFs. Next, we conducted additional ribosome profiling experiments on the 3T3-L1 preadipocyte cell line, chosen for its reliable differentiation into mature adipocytes upon induction with a cocktail containing insulin, 3-isobutyl-1-methylxanthine (IBMX), and dexamethasone. The differentiation process requires 7 days to advance from a pre-adipocyte to an adipocyte. To ensure we captured smORFs that might be transiently regulated during differentiation, we carried out ribosome profiling at three time points: undifferentiated, day 3 post-differentiation, and day 7 post-differentiation. Data from these experiments were processed through our established bioinformatic pipeline, which utilizes RibORF as the Ribo-Seq caller (11). The Ribo-Seq reads were mapped to the mm10 genome, resulting in 1,080 unannotated 3T3-L1 smORFs (Supplementary Table S1), of which 287 (∼27%) can be found within the SVF smORF dataset (Figure S1A). As we have seen in previous studies, the smORFs we identify overlap with the 5’-untranslated region (UTR), 3’-UTR, long non-coding RNAs, and intergenic regions. Of these, ∼62.2% of 3T3-L1 smORFs are those in the 5’-UTR, ∼19.5% in the 3’-UTR, and 15% overlap with non-coding RNAs and intergenic regions (Figure 2A). The median length of 3T3-L1 smORFs is 30 amino acids (Figure 2B), and the amino acid analysis shows a typical microprotein bias towards arginine, glycine, alanine, and tyrosine (Figure 2C). Here, we define an “unannotated smORF” as a smORF absent from GenBank and UniProt databases, identified by filtering against mouse reference protein sequences using BLASTp.

**Figure 2.**
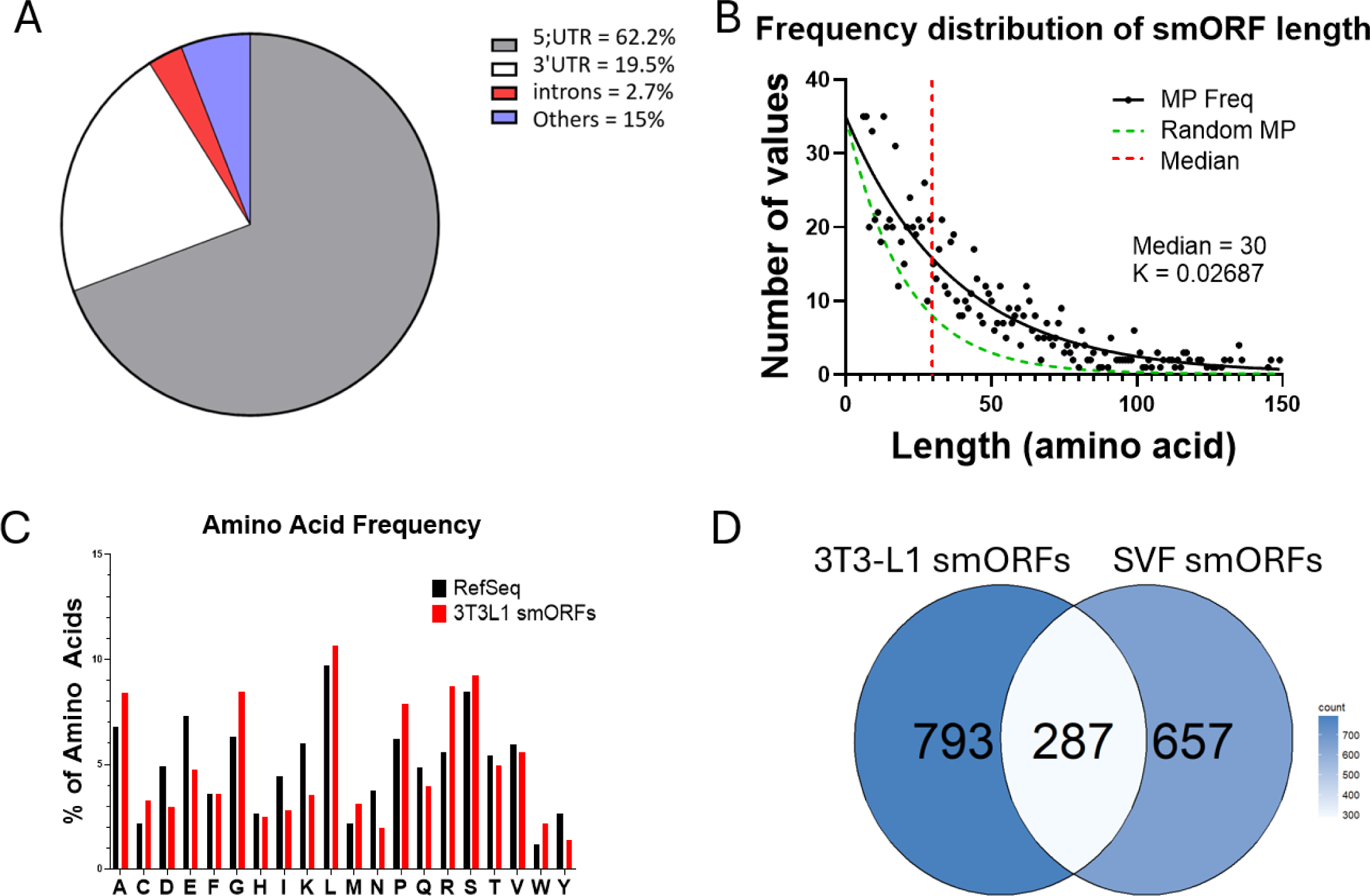
Identifying novel smORFs in 3T3-L1 cells by Ribosome Profiling. **A.** Pie chart showing smORF genomic location in the mm10 genome. Most smORFs overlap either 5’UTR, 3’UTR or intronic regions of the genome. **B.** Overall frequency distribution of smORFs identified from ribosome profiling. **C.** Amino acid frequency of predicted microprotein versus RefSeq (annotated genes). Letters refer to 1-letter amino acid codes. **D.** Venn diagram showing overlap of 3T3-L1 smORFs from this study versus SVF smORFs previously found in Martinez et al, 2023.

### A custom guide RNA library from 3T3-L1 and SVF smORFs and CRISPR Screen

For the unbiased analysis of smORF function in 3T3-L1 cells, we used a CRISPR/Cas9 screen. We generated our target library by combining the unannotated smORFs from the SVF smORFs and the newly acquired 3T3-L1 smORFs, which resulted in a total of 1737 unannotated Adipocyte-smORFs (Supplementary Table S1, Figure S1A). Since commercial guide RNA libraries do not include smORFs, we generated a custom guide RNA library using CCTop (34). Ideally, each smORF would be targeted by at least 10 guides, but one of the challenges with smORFs is the lack of genomic real estate found in longer genes. Nevertheless, we achieved adequate coverage with an average of ∼7 guides per smORF and 809 smORFs (∼48%) having 10 guides targeting their gene loci (Figure S1B). Note that 42 smORFs were excluded from the final smORF list due to the inability to design specific gRNAs to the target locus. The library also includes 192 annotated genes associated with cell proliferation and adipocyte function as positive controls for the screens and 250 non-targeting guide RNAs as negative controls, resulting in 12,925 guide RNAs. The guide RNAs were then cloned into a lentiviral library and infected into 3T3-L1 Cas9 cells at a multiplicity of infection (MOI) of <0.3 to ensure only one guide RNA per cell.

Identifying smORFs involved in 3T3-L1 adipocyte differentiation required us to distinguish smORFs essential for cell growth from smORFs that modulate differentiation. This was accomplished by allowing cells to proliferate for 30 days before adding the differentiation cocktail. smORFs involved in cell growth and proliferation could be delineated from those impacting differentiation by identifying guides that dropped out between day 0 and day 30. The remaining 3T3L1 cells were induced with a differentiation cocktail before sorting by FACS to identify smORFs that regulate adipocyte differentiation. The overall outline of our CRISPR screen can be found in Figure 1.

### smORFs that regulate cell proliferation

We compared the baseline cell population (immediately after lentiviral transduction of guide RNAs) with cells after 30 days (passage 10) using MaGECK MLE to identify smORFs affecting 3T3-L1 cell proliferation (35). The bioinformatics pipeline relied on CCTop for gRNA design and MaGeCK MLE to analyze the results from the subsequent CRISPR/Cas9 screen (Figure 3A). Results from three independent experiments are plotted with their corresponding B-scores from MaGECK with positive control genes highlighted in red, and smORFs in black (Figure 3B). Guide RNAs targeting genes essential for cell survival, such as mammalian target of rapamycin (mTOR), were nearly undetectable after 30 days in culture (Figure 3C), confirming that cells lacking mTOR lost proliferative capacity and dropped out of the guide RNA pool, validating the screen.

**Figure 3.**
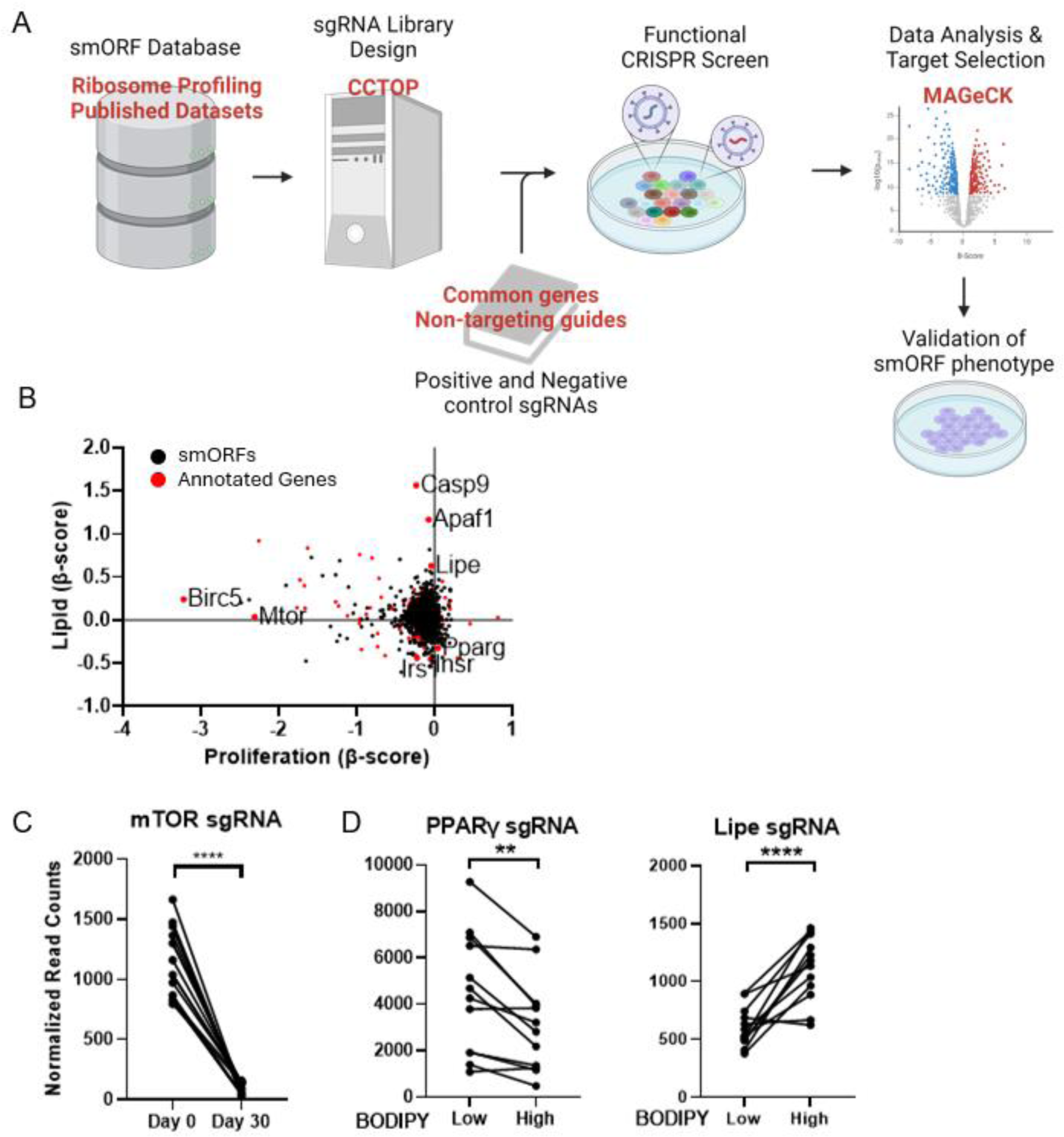
CRISPR-Cas9 screen of cell proliferation and lipid droplet size in 3T3-L1 cells **A.** Overall outline of the bioinformatic pipeline used for the CRISPR screen. Briefly, smORF genomic information, either from ribosome profiling or published datasets, is used by CCTOP to design specific sgRNAs. These sgRNAs are synthesized and cloned into lentiviral vectors for functional CRISPR/Cas9 screens. The cells are sequenced, and the data is analyzed by MAGeCK-MLE. SmORF candidates are validated through generation of knockout cell lines for downstream assays. **B.** Scatterplot of Proliferation vs Lipid β-scores calculated from MAGeCK-MLE. Annotated control genes are highlighted in Red, while smORFs are in Black. **C.** Normalized read counts of sgRNA targeting *mTOR* in 3T3-L1 Cas9 cells at Day 0 (immediately after sgRNA transduction) and Day 30 (after 10 passages). **D.** Normalized read counts of sgRNA targeting *PPARγ* and *Lipe* in BODIPY-low and BODIPY-high 3T3-L1 Cas9 cells. Statistics shown are Wald’s test calculated by MAGeCK-MLE. ** = P < 0.01; **** = p < 0.0001

To conservatively assess smORF phenotypes from the MaGECK MLE output, we applied a strict filter based on absolute β-score and FDR to generate a shortlist of candidates. For proliferation, we set an absolute β-score threshold of < 0.3, which aligns with genes like Cyclin D1 (Ccnd1; β-score = -0.32102) and Glucose-6-phosphatase catalytic subunit (G6pc; β-score = -0.30265) (Supplementary Table S2). This threshold yielded a shortlist of 109 smORFs with a significant phenotype on 3T3-L1 cell proliferation upon knockout (Supplementary Table S2). The adipocyte regulating smORFs were arbitrarily numbered from Adipocyte-smORF-1 to Adipocyte-smORF-1937 (see for Supplementary Table S1 for corresponding genomic coordinates). For example, Adipocyte-smORF-546 (β-score = -2.479), 543 (β-score = -1.906), 545 (β-score = -0.936), and 547 (β-score = -0.8657) have β-scores that are similar to genes such as mTOR (β-score = -2.306) and Hexokinase 2 (β-score = -0.8238) (Figure S2B-F). The smORFs listed above overlap with the 5’- and 3’- UTRs of the murine double minute 2 gene (Mdm2), a well-known negative regulator of P53 (Figure S2A), indicating that their role is more likely linked to the regulation of Mdm2 as smORFs instead of their independent actions as microproteins (36, 37).

### smORFs that regulate adipocyte biology

To identify smORFs that regulate adipocyte biology, we used the aforementioned FACS assay that stained differentiated 3T3-L1 cells with the lipid-staining dye BODIPY 493/503 (38), then used fluorescence-assisted cell sorting (FACS) to collect the top and bottom 5% of cells based on BODIPY signal intensity (Figure S3A). We note a recent study demonstrating that Cas9 overexpression in 3T3-L1 cells decreased adipogenesis (39). Furthermore, it is well documented that 3T3-L1 cells are susceptible to passage number, which may confound results due to varying cell culture conditions (40). We implemented several safeguards in our CRISPR/Cas9 screen to address these limitations. We repeated the experiment four times, using 3T3-L1-Cas9 cells at different passage numbers, and applied a conservative gating strategy during flow cytometry analysis (Figure S2B). Specifically, only the top and bottom 5% of BODIPY-labeled cells, representing the most and least differentiated cells, were collected in each experiment. Sequencing of the guide RNAs in these fractions and analysis by MaGECK MLE identifies those smORFs that can modulate lipid droplet formation during 3T3-L1 differentiation. This approach ensured the reliability and reproducibility of our findings.

Following FACS sorting, we were able to annotate guide RNAs that control lipid droplet formation in 3T3-L1 cells by sequencing. We observed a significant decrease in guide RNAs targeting Peroxisome Proliferator-Activated Receptor gamma (PPARγ) in BODIPY high cells compared to BODIPY low cells (Figure 3D), confirming that cells deficient in PPARγ have impaired adipocyte differentiation and cannot produce or store lipid droplets in differentiated 3T3-L1 cells. Conversely, we observed a significant increase in guide RNAs targeting Hormone Sensitive Lipase (Lipe) in BODIPY high cells, consistent with the role of Lipe as a rate-limiting enzyme in triglyceride breakdown in adipocytes; its knockout led to more triglyceride accumulation in 3T3-L1 cells, thus increasing BODIPY staining (Figure 3D). For lipid droplet accumulation, we similarly used an absolute β-score threshold of < 0.3, comparable to the effect range of PPARγ (β-score = -0.32699). This resulted in a shortlist of 38 smORFs that significantly affected lipid droplet accumulation in differentiated 3T3-L1 cells (Supplementary Table S2). Together, these data validate the robustness of our CRISPR/Cas9 screen and demonstrate its capability to identify novel smORF regulators of cell proliferation and adipocyte function.

### CRISPR screen does not show genomic location bias

Given that a CRISPR/Cas9 screen introduces genomic DNA mutations, we hypothesized that the genomic location of smORFs could impact screen outcomes. This hypothesis assumes that many predicted smORFs from our ribosome profiling experiments may be due to the impact of CRISPR/Cas9 targeting on the genome instead of the proteome. For example, in our Adipocyte smORF library of 1,695 predicted smORFs, 418 (∼24%) overlap with DNA repeat regions as defined by the RepeatMasker track on the UCSC Genome Browser. In this context, one hypothesis is that guide RNAs targeting smORFs overlapping repeat regions may induce multiple off-target mutations, potentially causing false positives. If true, a substantial proportion of smORF hits would be expected to overlap these repeat regions. However, among the 109 smORFs associated with a cell proliferation phenotype, only 24 (∼22%) overlap repeat regions (Figure 4A, B). Similarly, only six of the 38 smORFs with significant lipid droplet phenotypes (∼16%) overlap repeat regions (Figure 4A, 4B). Moreover, we observed no significant difference in β-score distribution between smORFs overlapping repeat regions and those in non-repetitive regions. These findings suggest that the inclusion of smORFs overlapping DNA repeat regions introduced minimal, if any, bias. However, this observational analysis does not rule out the possibility that some candidate smORF hits in repeat regions could be false positives, which would require targeted downstream validation.

**Figure 4.**
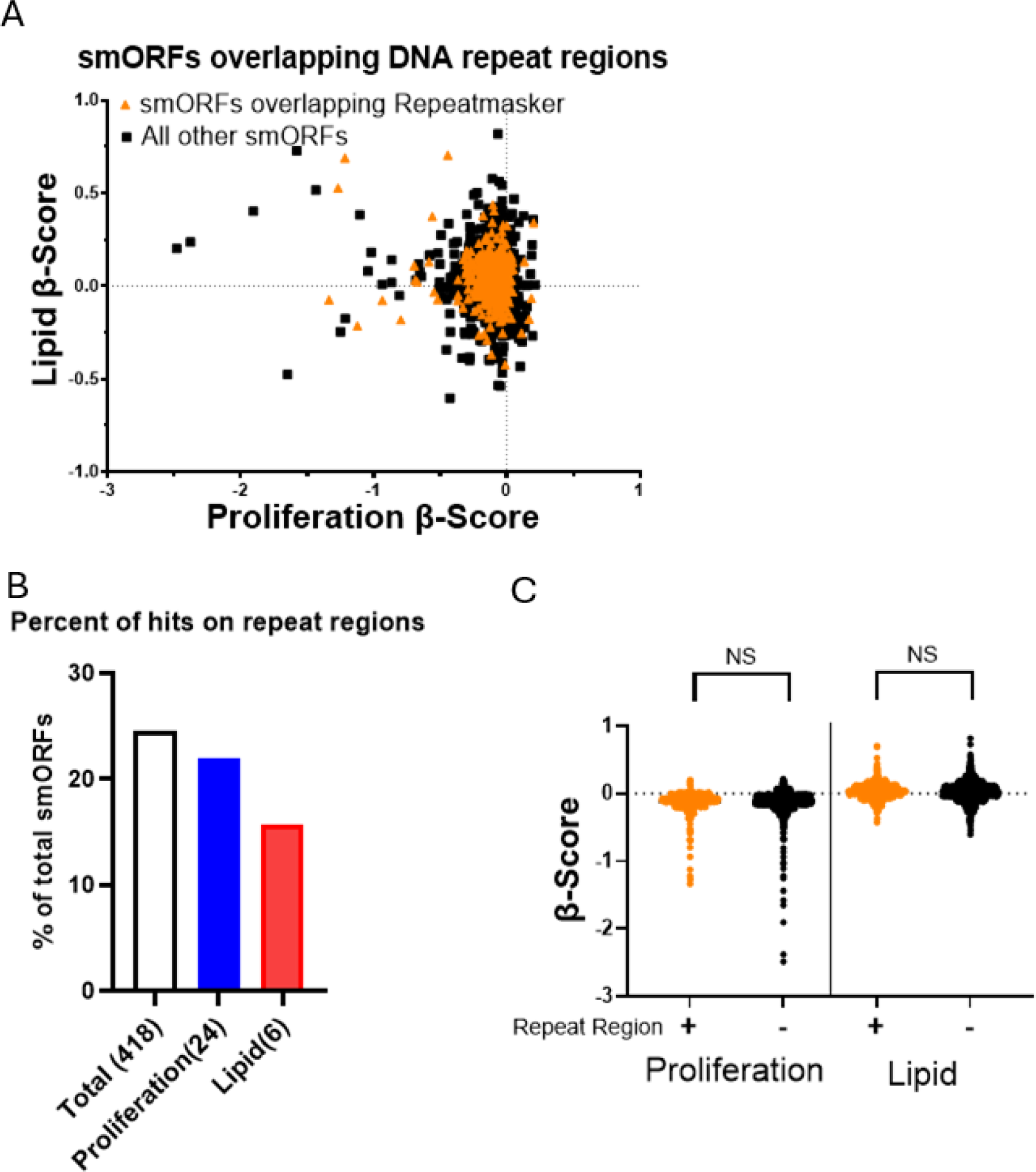
smORF location does not affect 3T3-L1 CRISPR screen outcome. **A.** Scatterplot of β-scores of smORFs overlapping DNA repeat regions (orange) versus other smORFs (black). β-scores were calculated from the MAGeCK-MLE. **B.** Bar graph showing percentage of smORFs overlapping DNA repeat regions from all smORFs, and smORFs that have significant proliferation or lipid droplet effects. Number of smORFs in each subgroup are in parenthesis. **C.** Distribution of β-scores from CRISPR/Cas9 Screen calculated by MAGeCK-MLE. No significant differences was observed, as calculated by Student’s t-Test.

An alternative hypothesis involving smORF location involves cis-regulation of 5’UTR and 3’UTR smORFs affecting the stability and function of the main transcript. For cell proliferation, our analysis showed a modest but significant decrease in β-scores for smORFs in 5’UTR or 3’UTR regions compared to smORFs in non-annotated regions (Figure S4A, S4B). Specifically, guide RNAs targeting 5’UTR or 3’UTR, on average, had weaker effects on cell proliferation than those targeting other smORF types (Figure S4B). However, as the difference in β-scores between categories is minor (less than 0.05), and thus no clear effect by genomic location. For lipid droplet accumulation, we observed no difference in β-scores between categories (Figure S4A), nor did we detect any significant deviation in smORF location between the entire smORF library and candidate smORF hits associated with proliferation or lipid droplet formation (Figure S4C). Taken together, these findings suggest the lack of a significant correlation between smORF genomic location and function.

### Analysis of smORF conservation versus functional outcome by CRISPR screen

While evolutionary conservation makes it more likely that a protein is functional, we wanted to explore the impact of conservation for smORFs and microproteins by examining the results from our unbiased screen. We assessed the conservation of our adipocyte smORF library across several species by running tBLASTn searches (using an E-value threshold of <0.01) against rat, chimpanzee, human, zebrafish, and Drosophila genomes (Figure 5A, Supplementary Table S3). Out of the 1,695 smORFs identified in mice, 778 (∼46%) had comparable sequences in the rat genome, while only 208 (∼12%) were similar in humans (Figure 5a). Further analysis of the β-scores from our CRISPR/Cas9 screen revealed no significant differences in cell proliferation or lipid droplet phenotypes between smORFs conserved in humans and those that were not (Figure 5B). Additionally, no enrichment of conserved human smORFs within the candidate smORFs showed effects on either proliferation or lipid droplet phenotypes (Figure 5C).

**Figure 5.**
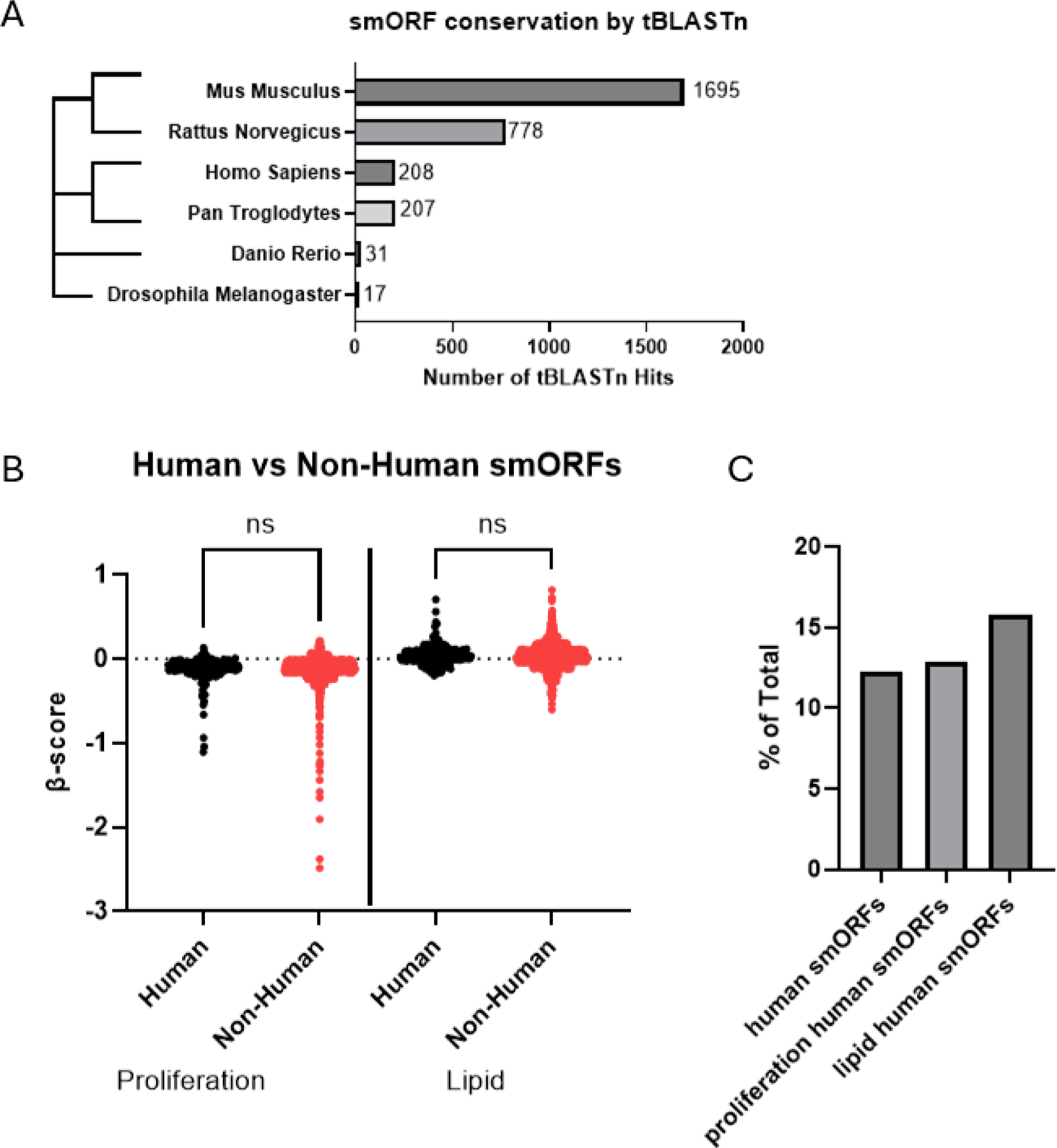
Conservation of smORFs does not correlate with CRISPR-Cas9 screen results. **A.** Unrooted phylogenic tree inferred from mitochondria DNA of closely related species. Bars indicate number of homologous sequences for novel smORFs that are found in the genome of represented species as predicted by tBLASTn. **B.** Distribution of β-scores from CRISPR/Cas9 Screen calculated by MAGeCK-MLE of smORFs that are conserved in human versus non-human smORFs. No significant differences were observed, as calculated by Student’s t-Test. **C.** Percentage of human smORFs cross different sub groups.

Here, we want to highlight two examples of smORFs that have varying degrees of conservation, as predicted by tBLASTn, with a significant effect on 3T3-L1 cell proliferation. The first is Adipocyte-smORF-543, which we highlighted earlier as having a significant anti-proliferative phenotype from the CRISPR screen (Figure S2A, S2D). Adipocyte-smORF-543 was predicted to be coded only within the rat genome by tBLASTn, albeit with a 10 AA deletion (Figure S5A). Another example is Adipocyte-smORF-340, which is predicted to be conserved human, chimp, rat, zebrafish, and drosophila by tBLASTn (Figure S5B). Adipocyte-smORF-340 encodes a 38 AA microprotein, which was shown to have a significant anti-proliferative effect in 7/10 sgRNA used within the CRISPR screen (Figure S5C). These differences highlight the growing appreciation for lineage-specific functional smORFs (41).

### Validating smORFs from Lipid Droplet Screen

To validate several of the smORF candidates influencing lipid droplet formation in differentiated 3T3-L1 cells, we generated individual knockout cell lines using CRISPR/Cas9, followed by differentiation into mature adipocytes and staining with Oil Red O to assess lipid droplet size. For our initial validation targets, we have chosen targets that match two criteria. First, as we are mainly interested in finding smORFs that are involved in adipocyte function, we excluded smORFs that exhibited a significant cell proliferation phenotype. Second, we double-checked the gRNA sequences used in the CRISPR/Cas9 screen to ensure the screen is robust (i.e., no off-target effects) and its effects are reproducible across experimental replicates. We specifically avoided checking for conservation across species to prioritize targets. Although conservation is a powerful indicator for smORF or microprotein function, we wanted to see if we could identify non-conserved yet functional microproteins as well. Representative images of three distinct knockout cell lines and a non-targeting control are shown in Figure 6A. The knockout of Adipocyte-smORF-1183 (1183 KO) in 3T3-L1 cells consistently led to a significant reduction in lipid droplet formation, as indicated by Oil Red O staining. Three of the four guide RNAs used for Adipocyte-smORF-1183 in the CRISPR/Cas9 screen were consistently enriched in the BODIPY-negative population across three separate experiments (Figure 6B).

**Figure 6.**
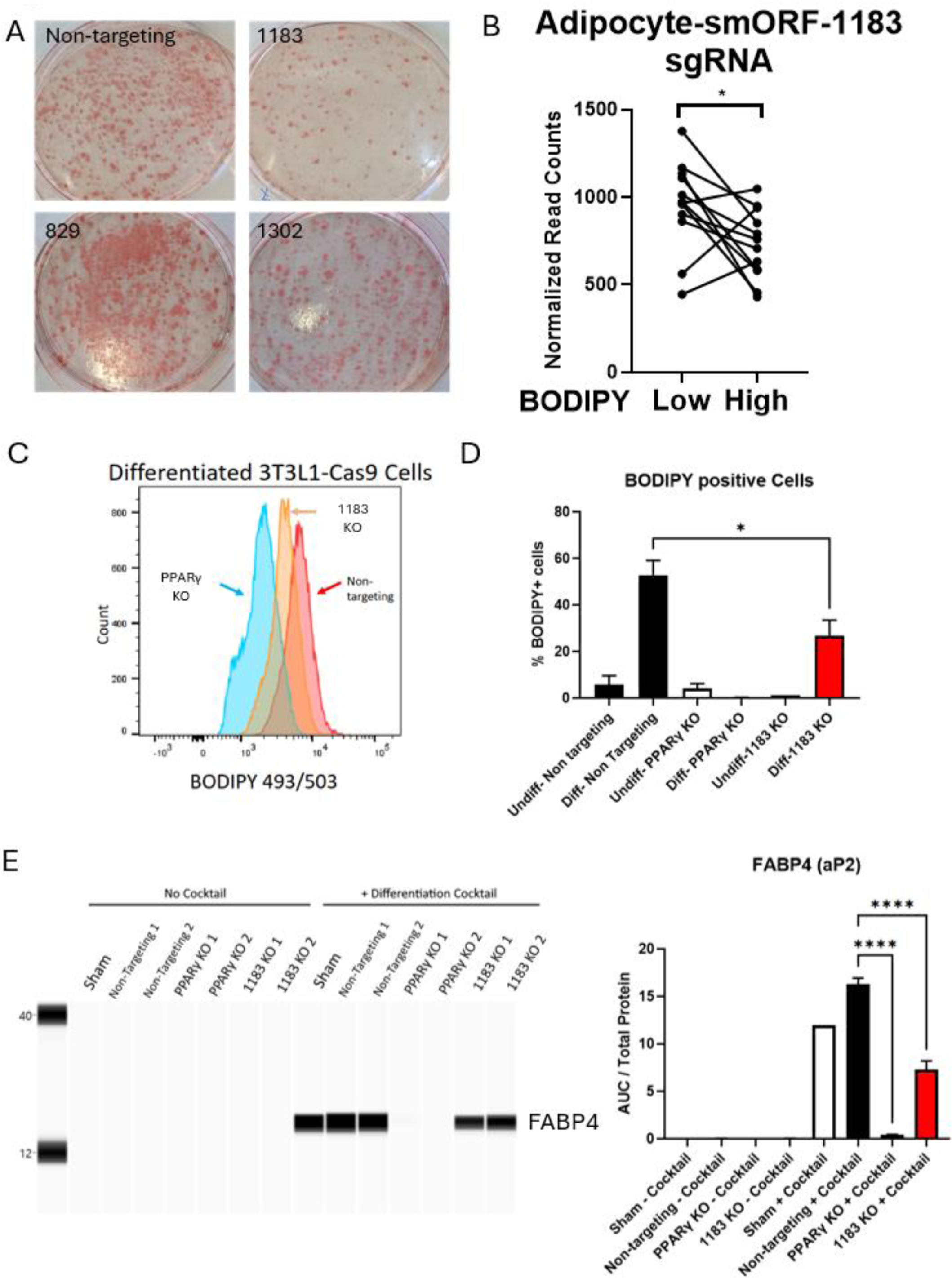
Validating the effect of Adipocyte-smORF-1183 on 3T3-L1 lipid droplet size. **A.** Representative images of Oil Red O stains of 3T3-L1 cas9 cells transduced with indicated sgRNAs. Images are captures on cells plated in 10cm plates. Numbers indicate specific Adipocyte-smORF 3T3-L1 knockout cell lines. **B.** Normalized read counts of Adipocyte-smORF-1183 sgRNA in the CRISPR/Cas9 screen of differentiated 3T3-L1 cells. P < 0.05 by Wald’s Test by MAGeCK-MLE. **C.** Flow cytometry analysis of differentiated 3T3-L1 Cas9 cells transduced with sgRNAs targeting either PPARγ(Blue), Adipocyte-smORF-1183 (Orange) or non-targeting control (Red). **D.** Bar graph showing percent of BODIPY positive cells quantified from three independent FACS experiments. Undiff = Undifferentiated, Diff = Differentiated 3T3-L1 cas9 cells transduced with indicated sgRNA. **E.** FABP4 levels of various conditions from capillary western blot (Protein Simple, Bio-Techne). Right panel shows quantification of measured spectrogram peaks (area under curve, AUC) Calculated statistics in panels D and E are by one-way ANOVA. * = P<0.05, **** = P< 0.0001

To verify this effect quantitatively, we analyzed BODIPY staining via flow cytometry in undifferentiated and differentiated 1183 KO cells (Figure 6C). Compared to PPARγ knockout cells (PPARγ KO) and cells with non-targeting control guides, 1183 KO cells showed an approximate 50% decrease in lipid droplet formation (Figure 6C, 6D). Additionally, analysis of FABP4 (Fatty Acid Binding Protein 4, also known as adipocyte protein 2)—a marker of mature adipocytes—demonstrated a significant reduction in protein expression in 1183 KO cells relative to controls in both undifferentiated and differentiated conditions (Figure 6E). Together, these results indicate that CRISPR/Cas9-mediated knockout of Adipocyte-smORF-1183 substantially impairs adipocyte lipid droplet formation.

### Validating the microprotein as the regulator of lipid droplets

From our bioinformatic analysis, Adipocyte-smORF-1183 is predicted to be encoded by exons 1, 3, and 4 of the annotated lncRNA 923011K14Rik (Figure S6A, S6B). As mentioned, one interesting aspect of these unbiased screens is the ability to find non-conserved microproteins with roles in biology. To this end, we focused on Adipocyte-smORF-1183 and set out to determine whether its encoded microprotein is regulating the lipid droplet phenotype. We cloned the Adipocyte-smORF-1183 into a mammalian expression vector and transfected it into 1183 KO cells, with cells transduced with non-targeting RNA as controls. Since these cells were generated through CRISPR/Cas9 knockout, we also designed mutants with alternative codons at the guide RNA binding site to reduce guide RNA targeting efficiency (Codon Replaced; CR mutants).

Additionally, we introduced methionine-to-alanine mutations at the first and eleventh amino acids to prevent translation of the microprotein, testing for the functionality of the non-coding RNA (M1A M11A mutants). This approach resulted in four distinct overexpression plasmids: wild-type (WT), WT;M1A M11A, CR, and CR;M1A M11A (Figure 7A). Using flow cytometry to quantify BODIPY staining in differentiated cells, we found that only WT and CR plasmids rescued the knockout phenotype in 1183 KO cells (Figure 7B). In contrast, the M1A M11A mutants for both WT and CR did not rescue the phenotype, indicating that translation of the microprotein is necessary for its function (Figure 7B).

**Figure 7.**
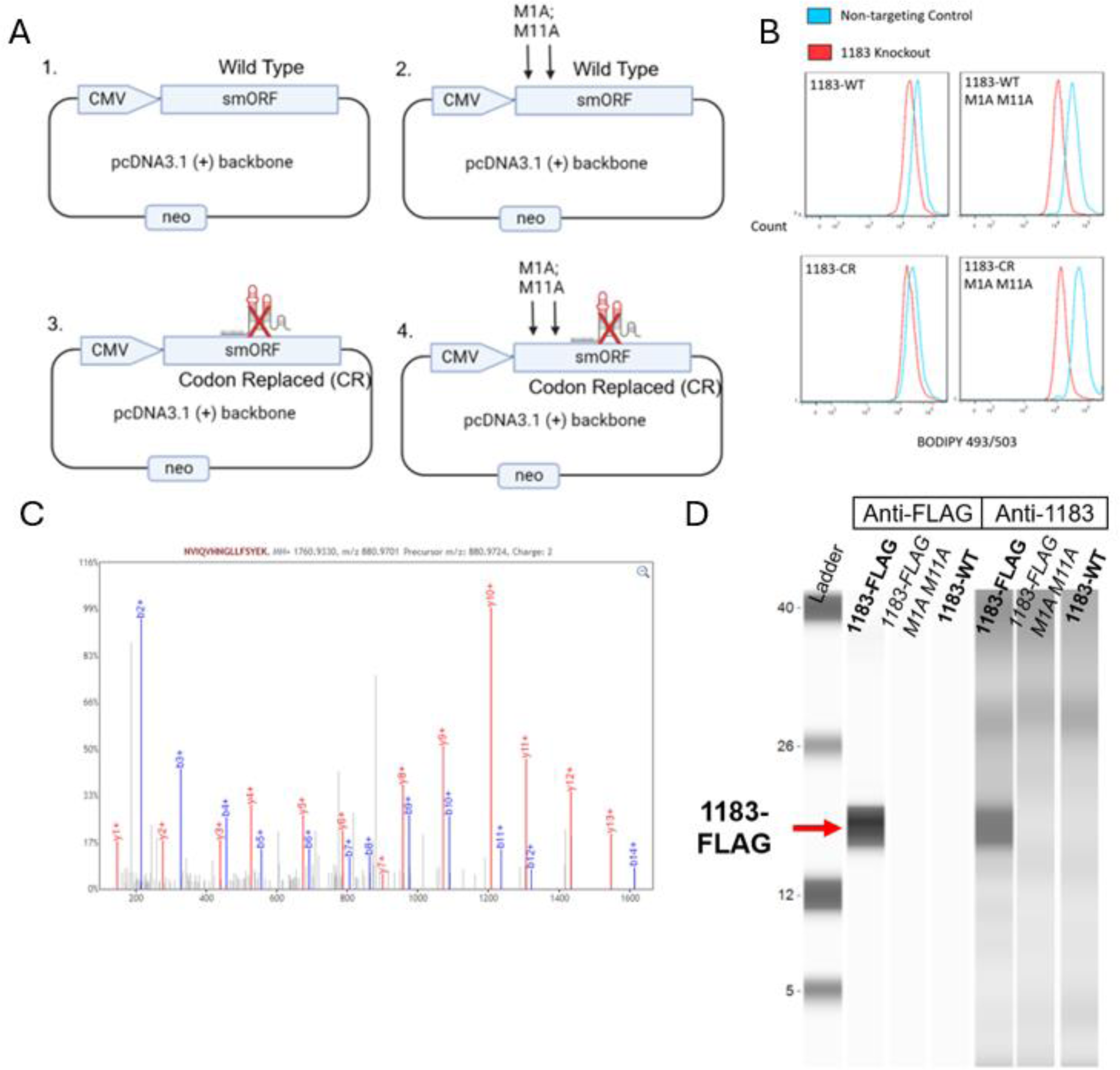
Overexpression of Adipocyte-smORF-1183 can rescue the 1183 KO phenotype**. A.** Overall experimental design of overexpression plasmids used in rescue experiments. Briefly, Adipocyte-smORF-1183 smORF is cloned into pcDNA3.1 (+) plasmid backbone downstream of the Cytomegalovirus promoter. **B.** Flow cytometry analysis of rescue experiment. Each panel indicates the overexpression plasmid transfected into either control cell line (blue) or Adipocyte-smORF-1183 knockout (red). **C.** Representative MS spectra of Adipocyte-smORF-1183 from HEK293T cell lysates transfected with wild type Adipocyte-smORF-1183. **D.** Capillary western blot of HEK293T cells transfected with indicated overexpression constructs.

To detect the microprotein, we overexpressed Adipocyte-smORF-1183-FLAG in HEK293T cells. We attempted to identify it using either western blotting (via the -FLAG epitope tag and the anti-Adipocyte-smORF-1183 antisera made in-house) or mass spectrometry. HEK293T cells were chosen due to the difficulty of transfecting 3T3-L1 cells and the need for large sample quantities for mass spectrometry. Adipocyte-smORF-1183 was detectable in HEK293T cells only when a FLAG epitope tag was added, as shown by both mass spectrometry and western blot (Figure 7C, 7D). We note that the native form of Adipocyte-smORF-1183 was undetectable by either method. We hypothesize that Adipocyte-smORF-1183 is a short-lived microprotein that influences lipid droplet formation in adipocytes or that a proteolytic processed form drives the biology (42, 43).

## Discussion

In this study, we leveraged CRISPR/Cas9 screening in the 3T3-L1 adipocyte differentiation model to identify microproteins influencing cell proliferation and lipid metabolism. Using an unbiased screen, we push the boundaries of microprotein research past the characterization of conserved microproteins and into the realm of lineage-specific smORFs. Through this approach, we identified and validated a novel microprotein, Adipocyte-smORF-1183, which significantly reduces lipid droplet formation during adipocyte differentiation. These findings highlight the importance of further investigating smORF functions across various biological systems and underscore the potential of CRISPR/Cas9 screens to uncover novel regulatory mechanisms (Figure 3A).

Our study demonstrates the importance of an unbiased approach in identifying functional smORFs using CRISPR/Cas9 screening. We found no significant differences in β-scores between smORFs that are conserved in the human genome and those that are not (Figure 5B). Additionally, human-conserved smORFs were not enriched among those with significant effects on cell proliferation or lipid droplet size after adipocyte differentiation (Figure 5C). These findings suggest that while conservation may serve as a valuable secondary criterion for prioritizing smORF validation, it should not be a primary determinant in initial screening efforts. The lack of bias towards conserved sequences likely results from a combination of factors, including the fact that the positions of many regulatory smORFs in 5’-UTRs predicted function as well as sequence in eukaryotes (44), and microproteins often favor intrinsic disorder (15, 45), where very short stretches of amino acids can dictate their function.

This approach enables the discovery of non-conserved smORFs with novel biological functions, such as Adipocyte-smORF-1183, which plays a key role in adipocyte lipid metabolism. Exploring species-specific smORFs like Adipocyte-smORF-1183 could lead to groundbreaking insights and the identification of new therapeutic targets for human diseases. Among the 109 smORFs identified as influencing cell proliferation, only 14 showed >50% conservation between human and mice. Likewise, of the 38 smORFs affecting lipid droplet metabolism in 3T3-L1 cells, we selected and characterized Adipocyte-smORF-1183, a rodent-specific microprotein, as an example of how lineage-specific smORFs can exert functional effects. If conservation had been used as a strict filtering criterion for our CRISPR/Cas9 screen, Adipocyte-smORF-1183 would not have been identified. While many functional smORFs are evolutionarily conserved, our findings suggest that conservation alone is not a reliable predictor of function. In the context of our smORF library, conservation did not significantly correlate with functional outcomes. Instead, our data reinforce the importance of experimental approaches in uncovering functional microproteins, particularly those that may have been overlooked due to lineage specificity.

Adipocyte-smORF-1183 is a predicted smORF whose gene location is within the RIKEN cDNA 9230114K14 gene (9230114K14Rik), previously annotated as a non-coding RNA (Figure S6). This validation provided additional confidence in our pipeline’s ability to identify functional microproteins influencing adipocyte differentiation. Rescue experiments in cells lacking Adipocyte-smORF-1183 confirmed that the encoded microprotein promotes lipid droplet formation in differentiated adipocytes, and knocking out Adipocyte-smORF-1183 resulted in a ∼50% reduction in lipid droplet accumulation in 3T3-L1 cells, further demonstrating its role in adipocyte biology.

Our CRISPR/Cas9 also provided a dataset for a systematic analysis of whether genomic/transcriptomic location can influence screening for functional smORFs. ORFs overlapping 5’ and 3’UTRs (uORFs and dORFs, respectively) have been shown to regulate the main ORF translation or mRNA translation, respectively, suggesting that uORFs or dORFs might only register as hits because of their role in regulating the translation of another protein instead of producing a functional microprotein (37). To assess whether traditional CRISPR/Cas9 screens can reveal such biases, we analyzed β-scores for smORFs based on their genomic categories. This concept was based on work that showed that CRISPR/Cas9 tiling across uORFs and ORFs can demonstrate distinct β-scores for uORFs and downstream ORFs (46).

For cell proliferation, we observed significant but minute differences between β-scores of 5’UTR and 3’UTR ORFs compared to smORFs in non-coding, intergenic, and other regions (Figure S4A, B). Non-coding, intergenic, and other smORFs had more potent effects on cell proliferation than those overlapping 5’UTR and 3’UTR regions. However, we found no significant differences in β-scores across smORF categories for lipid phenotypes. These results suggest that while potential cis-regulatory effects of uORFs and dORFs on main ORFs should be evaluated case-by-case during downstream validation, they are unlikely to dominate screen outcomes. Instead, non-coding, intergenic, and other smORFs appear to have more substantial effects on proliferation and may serve as primary targets for validation in CRISPR/Cas9 screens.

We also tested the hypothesis that smORFs overlapping repeat regions would lead to increased off-target effects. We found no significant differences in β-scores when comparing smORFs overlapping RepeatMasker tracks to those in unique genomic regions (Figure 4A). This suggests that the CCTop algorithm effectively minimizes off-target guide RNA design even for smORFs in repeat regions.

In summary, we demonstrate the efficacy of using CRISPR/Cas9 screens as a powerful method to identify functional smORFs from large ribosome profiling datasets. While prior studies employing this approach have primarily focused on cell proliferation, we extend its utility to interrogating complex phenotypes, such as adipocyte differentiation. This study establishes a robust framework for applying CRISPR/Cas9 screens to diverse biological contexts. Since our results identified a mouse-specific microprotein influencing adipocyte biology, conducting a similar study in human adipocytes could reveal human-specific microproteins with functional roles in adipose tissue regulation. Our screen results, performed on 3T3-L1 cells using a mouse adipocyte smORF library, will be a valuable resource for exploring smORF function in adipocyte biology.

## Materials and Methods

### Cell Culture

3T3-L1 (ATCC, CL-173) cell cultures used in the study are plated in 10 or 15cm plates with maintenance media: DMEM + 10% Bovine Calf Serum (Cytiva, SH30073.04). Cells are grown to ∼60-70% confluency before splitting 1:3 for general maintenance. For differentiating 3T3-L1 cells into adipocytes, cells are grown to confluency before changing into fresh maintenance media. After 24 hours, media is removed and replaced with differentiation media: DMEM + 10% FBS (Gibco, A5670701), 500uM 3-Isobutyl-1-methylxanthine (IBMX, Sigma, I5879), 5ug/ml Insulin (Sigma, I0516), and 1uM dexamethasone (Sigma, D1756). After 72 hours with differentiation media, cells are maintained in differentiation media without IBMX and dexamethasone for an additional 4-7 days with fresh media replaced every 48 hours.

### Preparation of 3T3-L1 for Ribosome Profiling and RNA-sequencing

For preparation of cells for RNA sequencing and Ribo-Seq, cells were seeded into 10cm plates at a density of 1 x 10^6^ cells per plate and allowed to grow to confluency before differentiation into mature adipocytes as described in protocol above. For both RNA-seq and Ribo-seq, the following experimental conditions were collected: day 0 (when cells are confluent), day 3 after induction of differentiation, and day 7 after induction of differentiation. For collection of RNAseq samples, cells were washed with PBS followed by addition of 500uL TRIZOL (Invitrogen Cat# 15596018) to each well and RNA is extracted following manufacturer’s protocol. The RNA samples were stored at -80°C until library prep and sequencing. For Ribo-seq samples, cells were washed twice after treatment with ice cold PBS + 100ug/mL cycloheximide (CHX; Fisher Scientific, AAJ66004X) and snap frozen in liquid nitrogen and stored at -80°C.

### Ribosome Profiling Library Preparation and sequencing

Library preparation for Ribosome Profiling was carried out as previously described (11). Briefly, cells were washed twice after treatment with ice cold PBS supplemented with 100ug/mL cycloheximide (CHX; Fisher Scientific, AAJ66004X) and snap frozen in liquid nitrogen and stored at -80°C. After thawing briefly on ice, the cells were lysed with 300uL of ice-cold lysis buffer (20mM Tris-HCl, pH 7.4, 150mM NaCl, 5mM MgCl2, 1% Triton X-100, 1mM DTT, 25U Turbo DNase (Thermofisher Cat# AM2238), and 100ug/mL CHX – added fresh) for 10 minutes with periodic vortexing and pipetting through an 18G needle to ensure the cells are dispersed. Cell lysates were clarified by centrifugation for 15,000g for 10 minutes at 4°C, flash frozen in liquid nitrogen, and stored in -80°C for up to 1 week. For each 3T3-L1 experimental condition, ribosome footprinting was carried out by digesting 50 mg of RNA in 300 mL lysate with 0.5 U/mg RNase I (Lucigen, N6901K) for 60 min at room temperature. Digestion reactions were quenched with 200 U Superase-In RNase I inhibitor (Thermo Fisher, AM2694) on ice. Following digestion, monosomes were purified from small RNA fragments using Micro-Spin S-400 HR columns (GE Life Sciences), and ribosome protected RNA fragments (RPFs) were extracted by acid phenol chloroform and isopropanol precipitation. Sequencing libraries were prepared as in McGlincy and Ingolia with some modifications (47). First, the RiboZero Mammalian Kit (Illumina) was used to deplete rRNA after RPF extraction and just prior to RPF size selection by gel extraction. Second, the Zymo clean & concentrator step after adaptor ligation is omitted and the reaction was carried over straight into reverse transcription. For the reverse transcription step to form cDNA, Episcript RT (Lucigen, ERT12910K) was used. Following reverse transcription, excess primer was degraded using Exonuclease I (Lucigen, X40520K) and the RNA templates were degraded using Hybridase (Lucigen, H39500). For the cDNA circularization step, CircLigase I (Lucigen, CL4111K) was used. PCR amplification was then carried out using Phusion Hot Start II High-Fidelity Master Mix (Thermo Fisher, F565L) for 9–12 cycles.

The adapters and primers for library construction used were as follows:

3’ adapter – 5’ /5phos/AGATCGGAAGAGCACACGTCTGAA/3ddC/-3’; RT primer – 5’-

/5Phos/AGATCGGAAGAGCGTCGTGTAGGGAAAGAG/iSp18/GTGACTGGAGTTCAGACGTGTGC C-3’; PCR forward primer – 5’-AATGATACGGCGACCACCGAGATCTACACTCTTTCCCTACACGACGCTC-3’ ; Illumina TruSeq Ribo Profile (Mammalian) Library Prep Kit index primers 1–12.

### RNA-Sequencing

Messenger RNA was purified from total RNA using poly-T oligo-attached magnetic beads. After fragmentation, the first strand cDNA was synthesized using random hexamer primers, followed by the second strand cDNA synthesis using either dUTP for directional library or dTTP for non-directional library. For the non-directional library, it was ready after end repair, A-tailing, adapter ligation, size selection, amplification, and purification. For the directional library, it was ready after end repair, A-tailing, adapter ligation, size selection, USER enzyme digestion, amplification, and purification. Library preparation used the Illumina Truseq stranded mRNA kit. The library was checked with Qubit and real-time PCR for quantification and bioanalyzer for size distribution detection. Quantified libraries were pooled and sequenced on NovaSeq6000, according to effective library concentration and data amount. The clustering of the index-coded samples was performed according to the manufacturer’s instructions. After cluster generation, the library preparations were sequenced on a NovaSeq6000 and paired-end reads were generated.

### Annotation of novel smORFs

The bioinformatic pipeline used to annotate smORFs is described in detail in Martinez et al with a few modifications (11). First, all reads were aligned to the mm10 reference genome using STAR v2.79a (48). Next, instead of Cufflinks (49), Stringtie v2.2.1 (50) was used to generate total transcriptome form RNA-seq reads, followed by 3-frame translation using a GTFtoFASTA (11). This generates a ORF database to which we used RibORF v0.1 (51, 52) to score the ribosome profiling reads. The ORF database was filtered using a threshold score of ≥ 0.7, ≤ 150 AA in length, and novel (not overlapping with known coding regions in RefSeq, and a maximum BLASTp alignment e-value to UniProt (53, 54)). The resulting smORF database was then annotated using Bedtools closest function against the mm10 reference genome to determine genome overlap (55). This also allowed splitting the smORFs into categories based on their genomic location. smORF overlaps between ribosome profiling reads were determined by quantifying read counts using HOMER analyzeRepeats and a count threshold of ≤ 10 was used (56). Conservation was determined using tBLASTn against the reference genomes of *Homo Sapiens, Pan troglodytes*, *Rattus norvegicus*, *Danio Rerio,* and *Drosophila Melanogaster* with a e-value threshold of ≤ 0.001 (57).

### Preparation of sgRNA library by CCTOP

To generate sgRNA for CRISPR library screen, we utilized CCTOP v1.0.1 (34). Briefly, Fasta file containing all smORF DNA sequences were compiled using *bedtools getfasta* (55), which is then fed into CCTOP (with the following parameters: *--pam NGG --fwdOverhang CACCG --revOverhang AAAC --maxOT 5*) to generate a large list of predicted sgRNAs with most smORFs. The generated list is then trimmed down to only contain a maximum number of 10 guides per smORF. For smORFs that failed to generate any sgRNA in this initial CCTOP run, we appended 10 nucleotides upstream and downstream of the DNA sequence and re-ran CCTOP. Sequences for both non-targeting sgRNA and known gene sgRNAs were retrieved from the published Brie library (58).

### Cloning of sgRNA library

The protocol clone to clone the sgRNA library into a lenti-viral backbone is described in detail in Joung et al (59) with a few modifications. First, the sgRNA sequence was appended with the following flanking sequences for initial PCR reaction as described in Wang et al (60).

5’ flanking sequence: TATCTTGTGGAAAGGACGAAACACCG

3’ flanking sequence: GTTTTAGAGCTAGAAATAGCAAGTTAAAAT.

The full pooled RNA library was produced by Twist Bioscience (San Francisco, CA) and PCR amplified as described in Joung et al before cloning into cloned into Lenti-Guide Puro (gift from Feng Zhang; Addgene plasmid # 52963). The amplified sgRNA was then transformed into Endura ElectroCompetent cells (Biosearch Tech, 60240) according to manufacturer’s instructions. For our library, a total of 5 electroporation reactions were done.

### Production of lentivirus

Production of either Cas9 lentivirus or sgRNA lentivirus was done by transfection of HEK293FT (Invitrogen, R70007) cells as described in Dull et al (61). Briefly, HEK293FT was maintained in DMEM + 10% FBS + 1% non-essential amino acid supplementation (Gibco, 11140050). For virus production, 1.7 x 10^7^ cells were plated on day zero. On Day one, a transfection mastermix was made containing the following 3mL of OptiMEM (Gibco, 31985062), 172.35ug of PEI-Max (Polysciences, 24765), 7.2ug of pMD2.G (gift from Didier Trono; Addgene plasmid # 12259), 13.25ug of pMDLg (gift from Didier Trono; Addgene plasmid # 12251), 5ug of pRSV-REV (gift from Didier Trono; Addgene plasmid # 12253), and either 20ug of cloned lenti-guide Puro, or 30 ug of lenti-Cas9-Blast (gift from Feng Zhang Addgene plasmid # 52962). The transfection mastermix was briefly vortexed for 30 seconds before incubation at room temperature for 10 minutes. Afterwards, 12mL of maintenance media was added to transfection mastermix, gently mixed, and added to HEK293FT cells, replacing the old media. The virus supernatant was collected 48 hours later. First, the supernatant is pooled into conical tubes and centrifuged for 10 minutes at 500g, 4°C to pellet any floating cells. Finally, the cleared supernatant is filtered through a 0.45 micron PES filter (Fisher Sci, 50-145-9633), aliquoted, and stored into -80C.

### Determining lentiviral titers

Lentiviral was transduced into 3T3L1 cells by spinfection and viral titers were determined by CellTiter-Glo (Promega, G7570) for cell viability after incubation with Blasticidin (Sigma, 15205) or Puromycin (Sigma, P8833) according to manufacturer’s description. Briefly, 3T3-L1 cells were collected to a concentration of 1 x 10^6^ cells/mL in standard media with an addition of 0.8ul/ml of polybrene (Sigma, TR-1003). Next, one million cells were added to each well of 12 well plates with varying volumes of virus supernatants of interest. The 12 well plate was spun for 1 hour at 1000g, 32°C before returning to the incubator for recovery. After 24 hours, the cells were replated into 96 well plates at 10,000 cells per well with addition of either Blasticidin (10ug/ml) or Puromycin (2ug/ml). Cell viability was measured with CellTiter-Glo after 72 hours with puromycin or after 5 days with blasticidin to determine viral titer. For downstream experiments involving Lenti-Cas9 Blast, we used a MOI of 0.7, and for Lenti-Guide Puro, we used a MOI of 0.3.

### CRISPR Library screen

For 3T3-L1 library screen, 3T3L1-Cas9 cells were first made by infecting Lenti-Cas9-Blast at a MOI of 0.7 by spinfection as described above. Next, a total of 300 million 3T3L1-Cas9 cells were transduced with the pooled sgRNA lentivirus by spinfection at a MOI of 0.3. After 24 hours, cells were replated in thirty 150mm plates at 10 million / plate and puromycin is added to select for transduced cells. Three days later, cells washed twice with PBS and pooled. Half of the pooled cells were snap frozen and stored in 80°C at this time as baseline day 0 cells. Next, 100 million cells were maintained and monitored to ensure <70% confluency before passaging every 3 days. After 9 passages (Around 27 days), cells were expanded into twenty plates. At passage 10, ten plates were collected and snap frozen as the day 30 experimental condition. The rest of the cells were differentiated into mature adipocytes as described above followed by FACS for lipid droplet size

### Lipid droplet staining, cell sorting, and flow cytometry analysis

Differentiated 3T3-L1 cells were stained for lipid droplets by BODIPY 493/503 (Invitrogen, D3922). Briefly, differentiated and non-differentiated 3T3-L1 cells were replaced fresh media containing 5mM BODIPY for 1 hour before washing with PBS and pooling in conical tubes. Cells were washed an additional 3 times in ice cold FACS buffer (PBS + 1% FBS) before resuspension in ice cold FACS buffer containing 200nM TO-PRO-3 (Invitrogen, T3605). Cells were filtered through 100micron and then 70 micron filters (Corning, CLS352350 & CLS352360) to achieve single cell resuspension before proceeding to FACS.

For FACS, the cells were maintained at 4C during the sorting process. Live cells were selected first by gating for TO-PRO-3 negative cells, followed by exclusion of debris using forward and side scatter pulse area parameters (FSC-A and SSC-A), exclusion of aggregates using pulse width (FSC-W and SSC-W), before gating populations based on BODIPY fluorescence. A BD Influx sorter and BD FACSAria Fusion sorter was used to sort cells, with PBS for sheath fluid (a 100-μm nozzle was used for cells with sheath pressure set to 20PSI). The cells were sorted directly into 50mL Falcon tubes containing media using the “1-drop Pure” sort mode. BODIPY 493/503 was excited with a 488nm laser (200mW for Influx, 100mW for Fusion) and detected using the “FITC” channel (530/40BP for Influx, 530/30BP for Fusion). TOPRO-3 was excited with a 640nm laser (120mW for Influx, 100mW for Fusion) and detected using the “APC” channel (670/30BP for both sorters). For analysis of various 3T3-L1 knockout cell lines, cells were analyzed by the BD LSRII using similar parameters and analyzed with FlowJo V10.

### DNA Extraction for sgRNA sequencing of CRISPR screen

The protocol for extracting DNA from 5 x 10^7^ cells is adapted from Chen et al (62). First, a lysis buffer consisting of 50mM Tris, pH8, 50mM EDTA, and 1% SDS is made. 6mL of lysis buffer is added to 5 x 10^7^ cell pellet with 30μl of proteinase K (20mg/mL, Qiagen, 19131) and incubated at 55°C overnight. Next day, 30 μl of RNase A (10mg/ml, Qiagen 19101) is added, mixed well by vortex, and incubated at 37°C for 30 minutes. The sample is cooled on ice for 5 minutes before adding 2mL of pre-chilled 7.5M ammonium acetate (Sigma A1542), mixed, and centrifuged at 4000g for 10 minutes at 4°C to precipitate the protein. The supernatant is carefully decanted into a new conical tube and re-centrifuged again at 4000g for 20 minutes at 4°C as an additional clean up step. Next, the supernatant is transferred to a new conical tube, and 6mL of 100% isopropanol (Sigma 190764) is added and mixed by inverting the tube repeatedly to precipitate the DNA. The DNA is spun down at 4000g for 10 minutes at 4°C and washed twice with 70% ethanol (Sigma, E7023) before airdrying and fully resuspending in TE buffer (Invitrogen, 12090015) by incubating at room temperature overnight.

For sgRNA sequencing, we followed the sgRNA sequencing library PCR protocol described in Joung et al with a maximum of 28 amplification cycles (59). Next up to 10 ug of PCR product is purified through gel extraction (Qiagen, 28704). The concentration and purity of the amplified library is confirmed by Qubit and Tapestation assays before sequencing on Illumina Nextseq 2000 with 20% Phi-X spike-in and single-end reads were generated.

### CRISPR Screen Data Analysis

For the CRISPR screen analysis we used MAGeCK v0.5.9.5 (35). Raw FASTQs from the sgRNA DNA sequencing were read by *mageck count* function, which generates count table for each sgRNA. The data analysis was done with *mageck mle* function with the following parameters: *--norm-method control – permutation-round 10 --remove-outliers.* This will generate an associated β-score matrix with both permutation and Wald’s Test P values. For our study, we generally only considered the Wald’s Test P-Values. Individual β-score plots were generated on Graphpad Prism, with the Wald’s test generated from MAGeCK analysis shown.

### Oil Red O staining of lipid droplets

The oil red O stock solution was made by first dissolving 0.35g of oil red O (Sigma, A12989.22) in 100mL of 100% isopropanol. This solution was then filtered through a 0.2 micron filter. The oil red O working solution was made by mixing 6mL of stock oil red O solution with 4mL of ddH2O. The working solution is incubated at room temperature for 20 minutes and filtered through 0.2 micron filter before use. Differentiated 3T3-L1 cells were washed with PBS, and fixed with 10% phosphate buffered formalin (Fisher Sci, SF100-4) for10 minutes. The fixative is then removed, and fresh formalin is added and incubated overnight. The next day, the fixative is removed, and cells were washed with ddH2O twice, followed by incubating in 60% isopropanol for 5 minutes. The cells were then let completely dry before stained with Oil Red O working solution for 10 minutes. After staining, the cells were washed with ddH2O four times before imaging.

### Protein sample preparation

For protein extraction, we used a general lysis buffer consisting of 50mM Tris pH 7.6, 150mM NaCl, 0.5% NP40, 5mM EDTA, 1mM sodium fluride, 0.1mM sodium orthovanadate, 1mM DTT, and 1x HALT protease inhibitor w/ EDTA (Thermofisher, 78429). Two ml lysis buffer was used per 10 cm plate, plates were scraped and lysates appropriately pooled. The lysates were centrifuged at 30 K xg for 30 min at 4C and the supernatants filtered through 5 mm syringe filters. The filtered samples were enriched for microproteins by binding to BondElut C8 cartridges, 40 mm sorbent, (Agilent Technologies, Santa Clara, CA). Approximately 100 mg sorbent was used per 10 mg total lysate protein. Cartridges were prepared with one column volume methanol and equilibrated with two volumes triethylammonium formate (TEAF) buffer pH 3 before the samples were applied. The cartridges were then washed with two column volumes of TEAF and the microprotein enriched fractions were eluted with acetonitrile:TEAF pH 3 (3:1) and lyophilized using a Speed-Vac concentrator. A Bradford protein assay (BioRad, Hercules, CA) was used to measure protein concentration of each sample after extraction and enrichment.

### Cloning and overexpression of Adipocyte-smORF-1183 in 3T3-L1 and HEK293T cells

Various overexpression plasmids of Adipocyte-smORF-1183 was cloned into pcDNA3.1(+) mammalian overexpression vector (Invitrogen) by Genscript (Piscataway, NJ, USA). The individual plasmids were amplified in TOP10 Chemically Competent E. Coli (Invitrogen, C404003) by Maxiprep (Qiagen, 12162). For overexpression of plasmids in 3T3-L1 cells, one hundred microliter of 4 x 10^6^ / mL cells were electroporated with 5.6ug of plasmid DNA per reaction according with NEON electroporator (Invitrogen MPK10096) according to manufacturer’s instructions with the following parameters: pulse voltage of 1300 volt, pulse width of 20ms, and pulse number of 2. After overexpression, cells were selected with Geneticin G418 (Gibco 10131035) at 50ug/mL for 1 week before experiments. For overexpression of Adipocyte-smORF-1183 in HEK293T, cells were transfected using lipofectamine 2000 according to manufacturer’s instructions.

### Antisera Production

#### Animal Care

All animal procedures were approved by the Institutional Animal Care and Use Committee of the Salk Institute and were conducted in accordance with the PHS Policy on Humane Care and Use of Laboratory Animals (PHS Policy, 2015), the U.S. Government Principles for Utilization and Care of Vertebrate Animals Used in Testing, Research and Training, the NRC Guide for Care and Use of Laboratory Animals (8th edition) and the USDA Animal Welfare Act and Regulations. Three 10 to 12-week old, specific pathogen free, female New Zealand white rabbits, weighing 3.0 to 3.2 kg at beginning of the study, were procured from Western Oregon Rabbit Co. (Philometh, OR) for use in antisera production. Rabbits were housed in an AAALAC accredited facility in a climate controlled environment (65-72 degrees Fahrenheit, 30-70% humidity) under 12-hour light/12-hour dark cycles. Rabbits were provided with ad libitum feed (5326 Lab Diet High Fiber), micro-filtered water and weekly fruits, vegetables and alfalfa hay for enrichment. Upon arrival, animals were physically examined by veterinary staff for good health and acclimated for at least two weeks prior to initiation of antiserum production. Each animal was monitored daily by the veterinary staff for signs of complications and weighed every two weeks. Routine physical exams were also performed by the veterinarian quarterly on all rabbits.

#### Preparation of Antigens

A synthetic peptide, Adipocyte-smORF-1183(55–82)-NH2 was synthesized by RS Synthesis (Louisville, KY), HPLC purified to >95%, and amino acid sequenced verified by mass spectrometry. The peptide was conjugated to maleimide activated Keyhole Limpet Hemocyanin (KLH) per manufacturer’s instructions (ThermoFisher) for use as immunogen.

The synthetic peptide sequence is CTQDQKRKQLDHLQLKNGDRNVIQVHNG-NH2.

#### Injection and Bleeding of Animals

The antigen was delivered to host animals using multiple intradermal injections of peptide-KLH conjugate in Complete Freund’s Adjuvant (initial inoculation) or incomplete Freund’s adjuvant (booster inoculations) every three weeks. Animals were bled, <10% total blood volume, one week following booster injections and bleeds screened for titer and specificity. Rabbits were administered 1-2 mg/kg Acepromazine IM prior to injections of antigen or blood withdrawal. At the termination of study, rabbits were exsanguinated under anesthesia (ketamine 50 mg/kg and aceprozamine 1 mg/kg, IM) and euthanized with an overdose of pentobarbital sodium and phenytoin sodium (1 ml/4.5 kg of body weight IC to effect). After blood was collected, the death of animals was confirmed. All animal procedures were conducted by experienced veterinary technicians, under the supervision of Salk Institute veterinarians.

#### Characterization of antisera

Each bleed from each rabbit was tested at multiple doses for the ability to recognize the synthetic peptide antigen; bleeds with highest titers were further analyzed by western immunoblot and/or JESS capillary western (Bio-Techne) for the ability to recognize the full-length endogenous protein and to check for cross-reactivity to other proteins. Serum from rabbit PBL #7559, with the best characteristics of titer against the synthetic peptide antigen, ability to recognize the endogenous protein, and specificity, was used for all studies.

### Capillary western blot

Capillary western blot for FABP4 (1:200, Abcam, ab92501) and FLAG epitope tag (1:200, cell signaling, 2368) was ran on the JESS (Bio-Techne) according to manufacturer’s instructions using 12-230 kda kit for FABP4 and 2-40 kda kit for microproteins. A protein concentration of 0.5mg/ml is used as initial loading material. Protein concentration was normalized using their internal protein standard for data analysis on the JESS software.

### Mass Spectrometry Analysis

C8 purified samples were precipitated by methanol/ chloroform and redissolved in 8 M urea/100 mM TEAB, pH 8.5. Proteins were reduced with 5 mM tris(2-carboxyethyl)phosphine hydrochloride (TCEP, Sigma-Aldrich) and alkylated with 10 mM chloroacetamide (Sigma-Aldrich). Proteins were digested overnight at 37 °C in 2 M urea/100 mM TEAB, pH 8.5, with trypsin (Promega). Digestion was quenched with formic acid, 5 % final concentration.

Digested samples were run on a Thermo Q Exactive. The samples were injected directly onto a 25 cm, 100 μm ID column packed with BEH 1.7 μm C18 resin. Samples were separated at a flow rate of 300 nl/min on a nLC 1000. Buffer A and B were 0.1% formic acid in water and 0.1% formic acid in acetonitrile, respectively. A gradient of 5–25% B over 280 min, an increase to 40% B over 60 min, an increase to 90% B over another 10 min and a hold at 90%B for a final 10 min was used for a total run time of 360 min. The column was re-equilibrated with 15 μl of buffer A prior to the injection of sample. Peptides were eluted directly from the tip of the column and nanosprayed into the mass spectrometer by application of 2.8 kV voltage at the back of the column. The Q Exactive was operated in data dependent mode. Full MS1 scans were collected in the Orbitrap at 70K resolution with a mass range of 400 to 1800 m/z and an AGC target of 5e^6^. A top 10 method was used for MS/MS with an AGC target of 5e^6^, resolution of 17.5K, minimum intensity of 5000 and NCE of 25. Maximum fill times were set to 120 ms and 500 ms for MS and MS/MS scans, respectively. Quadrupole isolation at 2 m/z was used, monoisotopic precursor selection was enabled, charge states of 2–7 were selected and dynamic exclusion was used with an exclusion duration of 15 s.

Protein and peptide identification were done with Integrated Proteomics Pipeline – IP2 (Integrated Proteomics Applications). Tandem mass spectra were extracted from raw files using RawConverter (63) and searched with ProLuCID (64) against a nonredundant mouse Uniprot database appended with adipocyte smORF sequences. The search space included all fully-tryptic and half-tryptic peptide candidates. Carbamidomethylation on cysteine was considered as a static modification. Data was searched with 50 ppm precursor ion tolerance and 600 ppm fragment ion tolerance. Identified proteins were filtered to using DTASelect (65) and utilizing a target-decoy database search strategy to control the false discovery rate to 1% at the protein level (66). A minimum of 1 peptide per protein and 1 tryptic end per peptide were required and precursor delta mass cutoff of 10ppm.

## Acknowledgments

The authors would like to thank the NIH (F32 DK132927, RC2 DK129961, R01 DK106210, R01 GM102491 and RF1 AG086547), Ferring Foundation, and Clayton Foundation for support of their work. The Flow Cytometry Core Facility at Salk Institute is supported by the Salk NIH-NCI Cancer Center P30 014195 grant, NIH Shared Instrumentation Grants S10-OD023689 and S10-OD034268, with additional funding support from Larry and Carol Greenfield Technology Fund.

## References

1. M. A. Basrai, P. Hieter, J. D. Boeke, Small open reading frames: beautiful needles in the haystack. Genome Res 7, 768–771 (1997).

2. J. P. Couso, P. Patraquim, Classification and function of small open reading frames. Nat Rev Mol Cell Biol 18, 575–589 (2017).

3. J. Ruiz-Orera, M. M. Alba, Translation of Small Open Reading Frames: Roles in Regulation and Evolutionary Innovation. Trends Genet 35, 186–198 (2019).

4. A. Saghatelian, J. P. Couso, Discovery and characterization of smORF-encoded bioactive polypeptides. Nat Chem Biol 11, 909–916 (2015).

5. G. Storz, Y. I. Wolf, K. S. Ramamurthi, Small proteins can no longer be ignored. Annu Rev Biochem 83, 753–777 (2014).

6. J. Lawrence, When ELFs are ORFs, but don’t act like them. Trends Genet 19, 131–132 (2003).

7. H. Ochman, Distinguishing the ORFs from the ELFs: short bacterial genes and the annotation of genomes. Trends Genet 18, 335–337 (2002).

8. J. J. Mohsen, A. A. Martel, S. A. Slavoff, Microproteins-Discovery, structure, and function. Proteomics 23, e2100211 (2023).

9. N. T. Ingolia et al., Ribosome profiling reveals pervasive translation outside of annotated protein-coding genes. Cell Rep 8, 1365–1379 (2014).

10. N. T. Ingolia, S. Ghaemmaghami, J. R. Newman, J. S. Weissman, Genome-wide analysis in vivo of translation with nucleotide resolution using ribosome profiling. Science 324, 218–223 (2009).

11. T. F. Martinez et al., Accurate annotation of human protein-coding small open reading frames. Nat Chem Biol 16, 458–468 (2020).

12. T. F. Martinez et al., Profiling mouse brown and white adipocytes to identify metabolically relevant small ORFs and functional microproteins. Cell Metab 35, 166–183.e111 (2023).

13. J. M. Mudge et al., Standardized annotation of translated open reading frames. Nat Biotechnol 40, 994–999 (2022).

14. B. R. Nelson et al., A peptide encoded by a transcript annotated as long noncoding RNA enhances SERCA activity in muscle. Science 351, 271–275 (2016).

15. N. Arnoult et al., Regulation of DNA repair pathway choice in S and G2 phases by the NHEJ inhibitor CYREN. Nature 549, 548–552 (2017).

16. P. J. Hung et al., MRI Is a DNA Damage Response Adaptor during Classical Non-homologous End Joining. Mol Cell 71, 332–342 e338 (2018).

17. C. Liang et al., Mitochondrial microproteins link metabolic cues to respiratory chain biogenesis. Cell Rep 40, 111204 (2022).

18. S. Zhang et al., LINC00116-encoded microprotein mitoregulin regulates fatty acid metabolism at the mitochondrial outer membrane. iScience 26, 107558 (2023).

19. S. Zhang et al., Mitochondrial peptide BRAWNIN is essential for vertebrate respiratory complex III assembly. Nat Commun 11, 1312 (2020).

20. Y. Chen et al., The microprotein HDSP promotes gastric cancer progression through activating the MECOM-SPINK1-EGFR signaling axis. Nat Commun 15, 8381 (2024).

21. M. Polycarpou-Schwarz et al., The cancer-associated microprotein CASIMO1 controls cell proliferation and interacts with squalene epoxidase modulating lipid droplet formation. Oncogene 37, 4750–4768 (2018).

22. J. E. Yang et al., LINC00998-encoded micropeptide SMIM30 promotes the G1/S transition of cell cycle by regulating cytosolic calcium level. Mol Oncol 17, 901–916 (2023).

23. P. Poirier et al., Obesity and cardiovascular disease: pathophysiology, evaluation, and effect of weight loss: an update of the 1997 American Heart Association Scientific Statement on Obesity and Heart Disease from the Obesity Committee of the Council on Nutrition, Physical Activity, and Metabolism. Circulation 113, 898–918 (2006).

24. T. M. Powell-Wiley et al., Obesity and Cardiovascular Disease: A Scientific Statement From the American Heart Association. Circulation 143, e984–e1010 (2021).

25. K. D. Hall, S. Kahan, Maintenance of Lost Weight and Long-Term Management of Obesity. Med Clin North Am 102, 183–197 (2018).

26. R. W. Jeffery et al., Long-term maintenance of weight loss: current status. Health Psychol 19, 5–16 (2000).

27. J. G. Kang, C. Y. Park, Anti-Obesity Drugs: A Review about Their Effects and Safety. Diabetes Metab J 36, 13–25 (2012).

28. A. M. Jastreboff et al., Tirzepatide Once Weekly for the Treatment of Obesity. N Engl J Med 387, 205–216 (2022).

29. Q. Chu et al., Regulation of the ER stress response by a mitochondrial microprotein. Nat Commun 10, 4883 (2019).

30. A. L. Rocha et al., An Inner Mitochondrial Membrane Microprotein from the SLC35A4 Upstream ORF Regulates Cellular Metabolism. J Mol Biol 436, 168559 (2024).

31. J. Chen et al., Pervasive functional translation of noncanonical human open reading frames. Science 367, 1140–1146 (2020).

32. E. Xu, J. Zhang, X. Chen, MDM2 expression is repressed by the RNA-binding protein RNPC1 via mRNA stability. Oncogene 32, 2169–2178 (2013).

33. J. S. Mattick et al., Long non-coding RNAs: definitions, functions, challenges and recommendations. Nat Rev Mol Cell Biol 24, 430–447 (2023).

34. M. Stemmer, T. Thumberger, M. Del Sol Keyer, J. Wittbrodt, J. L. Mateo, CCTop: An Intuitive, Flexible and Reliable CRISPR/Cas9 Target Prediction Tool. PLoS One 10, e0124633 (2015).

35. W. Li et al., MAGeCK enables robust identification of essential genes from genome-scale CRISPR/Cas9 knockout screens. Genome Biol 15, 554 (2014).

36. D. A. Freedman, L. Wu, A. J. Levine, Functions of the MDM2 oncoprotein. Cell Mol Life Sci 55, 96–107 (1999).

37. Q. Wu et al., Translation of small downstream ORFs enhances translation of canonical main open reading frames. Embo j 39, e104763 (2020).

38. C. Durandt et al., Novel flow cytometric approach for the detection of adipocyte subpopulations during adipogenesis. J Lipid Res 57, 729–742 (2016).

39. T. Suchý et al., Evaluating the feasibility of Cas9 overexpression in 3T3-L1 cells for generation of genetic knock-out adipocyte cell lines. Adipocyte 10, 631–645 (2021).

40. E. Cave, N. J. Crowther, The Use of 3T3-L1 Murine Preadipocytes as a Model of Adipogenesis. Methods Mol Biol 1916, 263–272 (2019).

41. N. Vakirlis, Z. Vance, K. M. Duggan, A. McLysaght, De novo birth of functional microproteins in the human lineage. Cell Rep 41, 111808 (2022).

42. W. T. Liu et al., Imaging mass spectrometry of intraspecies metabolic exchange revealed the cannibalistic factors of Bacillus subtilis. Proc Natl Acad Sci U S A 107, 16286–16290 (2010).

43. C. S. Stein et al., Mitoregulin: A lncRNA-Encoded Microprotein that Supports Mitochondrial Supercomplexes and Respiratory Efficiency. Cell Rep 23, 3710–3720.e3718 (2018).

44. G. E. May et al., Unraveling the influences of sequence and position on yeast uORF activity using massively parallel reporter systems and machine learning. Elife 12 (2023).

45. N. G. D’Lima et al., A human microprotein that interacts with the mRNA decapping complex. Nat Chem Biol 13, 174–180 (2017).

46. J. R. Prensner et al., Noncanonical open reading frames encode functional proteins essential for cancer cell survival. Nat Biotechnol 39, 697–704 (2021).

47. N. J. McGlincy, N. T. Ingolia, Transcriptome-wide measurement of translation by ribosome profiling. Methods 126, 112–129 (2017).

48. A. Dobin et al., STAR: ultrafast universal RNA-seq aligner. Bioinformatics 29, 15–21 (2013).

49. C. Trapnell et al., Differential gene and transcript expression analysis of RNA-seq experiments with TopHat and Cufflinks. Nat Protoc 7, 562–578 (2012).

50. M. Pertea et al., StringTie enables improved reconstruction of a transcriptome from RNA-seq reads. Nat Biotechnol 33, 290–295 (2015).

51. Z. Ji, RibORF: Identifying Genome-Wide Translated Open Reading Frames Using Ribosome Profiling. Curr Protoc Mol Biol 124, e67 (2018).

52. Z. Ji, R. Song, A. Regev, K. Struhl, Many lncRNAs, 5’UTRs, and pseudogenes are translated and some are likely to express functional proteins. Elife 4, e08890 (2015).

53. Anonymous, UniProt: the Universal Protein Knowledgebase in 2023. Nucleic Acids Res 51, D523–d531 (2023).

54. A. Bairoch, R. Apweiler, The SWISS-PROT protein sequence database: its relevance to human molecular medical research. J Mol Med (Berl) 75, 312–316 (1997).

55. A. R. Quinlan, I. M. Hall, BEDTools: a flexible suite of utilities for comparing genomic features. Bioinformatics 26, 841–842 (2010).

56. S. Heinz et al., Simple combinations of lineage-determining transcription factors prime cis-regulatory elements required for macrophage and B cell identities. Mol Cell 38, 576–589 (2010).

57. C. Camacho et al., BLAST+: architecture and applications. BMC Bioinformatics 10, 421 (2009).

58. J. G. Doench et al., Optimized sgRNA design to maximize activity and minimize off-target effects of CRISPR-Cas9. Nat Biotechnol 34, 184–191 (2016).

59. J. Joung et al., Genome-scale CRISPR-Cas9 knockout and transcriptional activation screening. Nat Protoc 12, 828–863 (2017).

60. T. Wang, E. S. Lander, D. M. Sabatini, Single Guide RNA Library Design and Construction. Cold Spring Harb Protoc 2016, pdb.prot090803 (2016).

61. T. Dull et al., A third-generation lentivirus vector with a conditional packaging system. J Virol 72, 8463–8471 (1998).

62. S. Chen et al., Genome-wide CRISPR screen in a mouse model of tumor growth and metastasis. Cell 160, 1246–1260 (2015).

63. L. He, J. Diedrich, Y. Y. Chu, J. R. Yates, 3rd, Extracting Accurate Precursor Information for Tandem Mass Spectra by RawConverter. Anal Chem 87, 11361–11367 (2015).

64. T. Xu et al., ProLuCID: An improved SEQUEST-like algorithm with enhanced sensitivity and specificity. J Proteomics 129, 16–24 (2015).

65. D. L. Tabb, W. H. McDonald, J. R. Yates, 3rd, DTASelect and Contrast: tools for assembling and comparing protein identifications from shotgun proteomics. J Proteome Res 1, 21–26 (2002).

66. J. Peng, J. E. Elias, C. C. Thoreen, L. J. Licklider, S. P. Gygi, Evaluation of multidimensional chromatography coupled with tandem mass spectrometry (LC/LC-MS/MS) for large-scale protein analysis: the yeast proteome. J Proteome Res 2, 43–50 (2003).

