## Supplementary File for "CRISPR-Cas9 Screening Reveals Microproteins Regulating Adipocyte Proliferation and Lipid Metabolism"

**This PDF file includes:**

Figures S1 to S6

Legends for Supplementary Tables S1 to S3

**Figures**

Figure S1

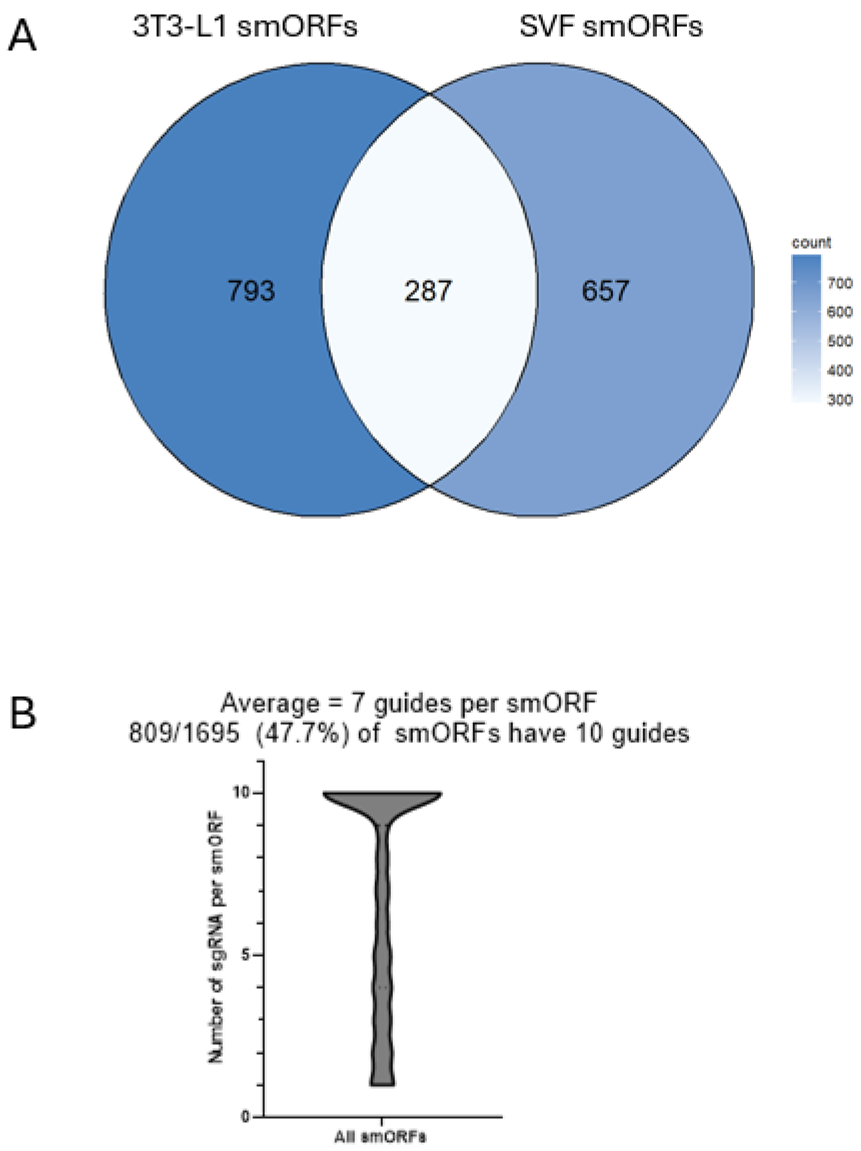


**Figure S1**. Description of smORFs and sgRNA used in the CRISPR/Cas9 screen**. A**. Venn diagram showing source of ribosome profiling data to generate list of smORF used. 3T3-L1 smORFs are from this experiment, and SVF smORFs were annotated from Martinez et al, 2023**. B.** Graph showing distribution of the number of gRNAs designed for all smORFs.

**Figure S2**

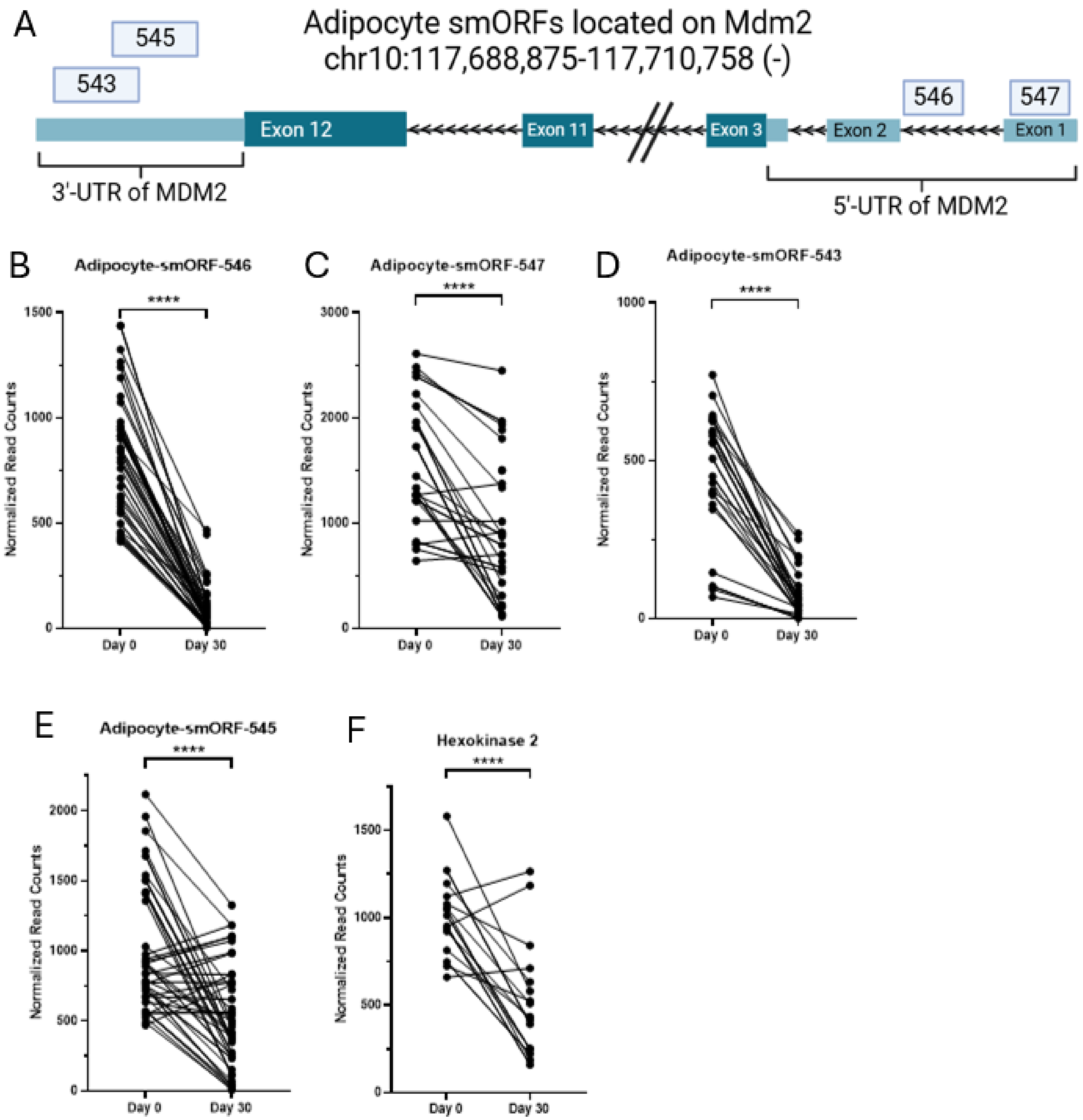


**Figure S2.** Adipocyte smORFs that overlap the 5' and 3' UTR of MDM2. **A.** Diagram showing the location of the 4 adipocyte smORFs that overlap the Mdm2 gene. **B-F.** Normalized read counts of sgRNA targeting various smORFs and Hexokinase 2 in 3T3-L1 Cas9 cells at Day 0 (immediately after sgRNA transduction) and Day 30 (after 10 passages). **** = P < 0.0001, by Wald's Test as calculated by MAGeCK-MLE.

**Figure S3**

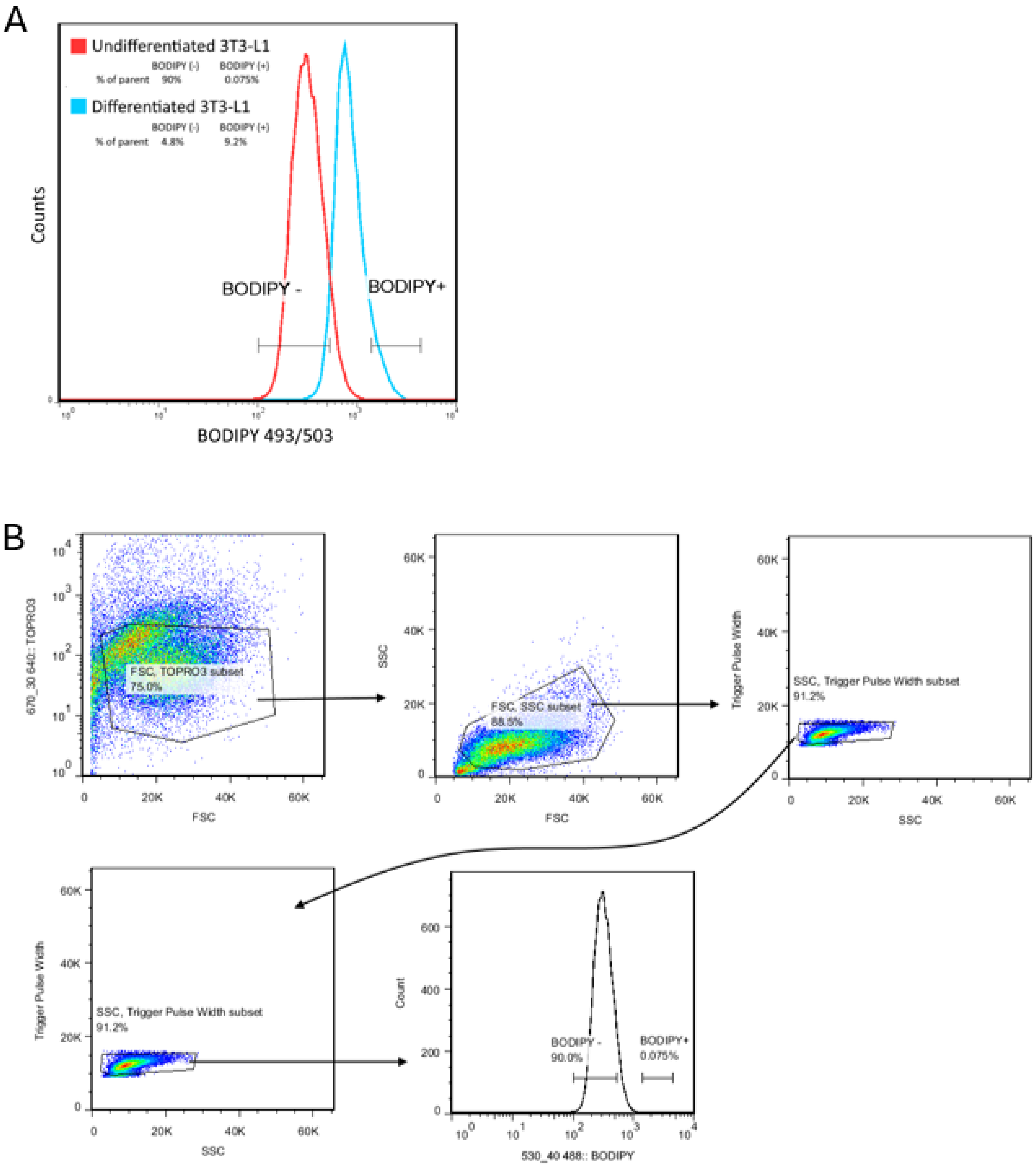


**Figure S3.** Overview of flow cytometry analysis of BODIPY in differentiated 3T3-L1 adipocytes**. A.** Example of gating used for FACS collection of differentiated 3T3-L1 Cas9 cells in the CRISPR-Cas9 library screen. In general, a gating of < 10% of the cell population was used for BODIPY positive cell population**. B.** Overall gating strategy used for flow cytometry analysis. Cells were gated first for live/dead (TO-PRO3), followed by forward and side scatter gates for singletons before gating for BODIPY.

### Figure S4

A

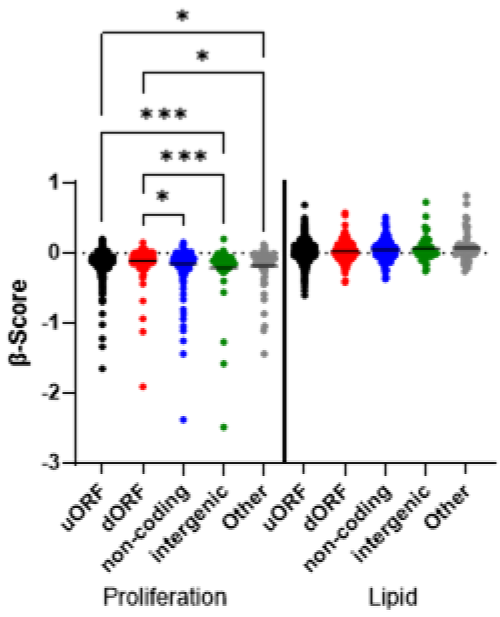


B

| | uORF | dORF | non-coding | intergenic | Other |
|---|---|---|---|---|---|
| Proliferation | -0.1197 ± 0.1344 | -0.1067 ± 0.1744 | -0.1557 ± 0.259 | -0.2118 ± 0.405 | -0.1852 ± 0.2795 |
| Lipid | 0.0259 ± 0.1326 | 0.02799 ± 0.1283 | 0.0465 ± 0.1385 | 0.06377 ± 0.1638 | 0.07744 ± 0.2037 |

C

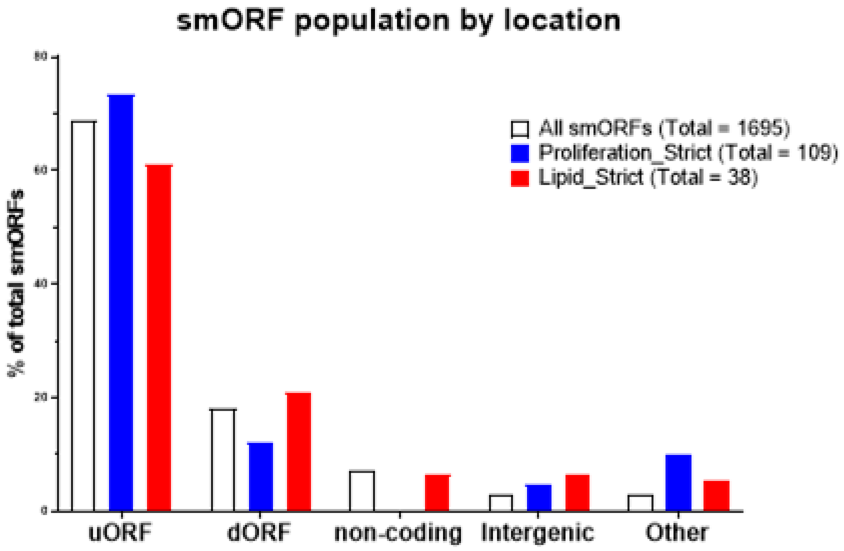


**Figure S4.** smORF genomic location and CRISPR-Cas9 screen outcome. **A.** Distribution of β-scores calculated from 3T3-L1 screens by MAGeCK-MLE. Statistics shown are calculated by one-way ANOVA only within proliferation & lipid subgroups. * = P<0.05, *** = P < 0.001. **B.** Table showing calculated average ±

standard deviation of β-scores as shown in Figure S4A. **C.** Bar graph showing distribution of smORFs among different genomic locations.

**Figure S5**

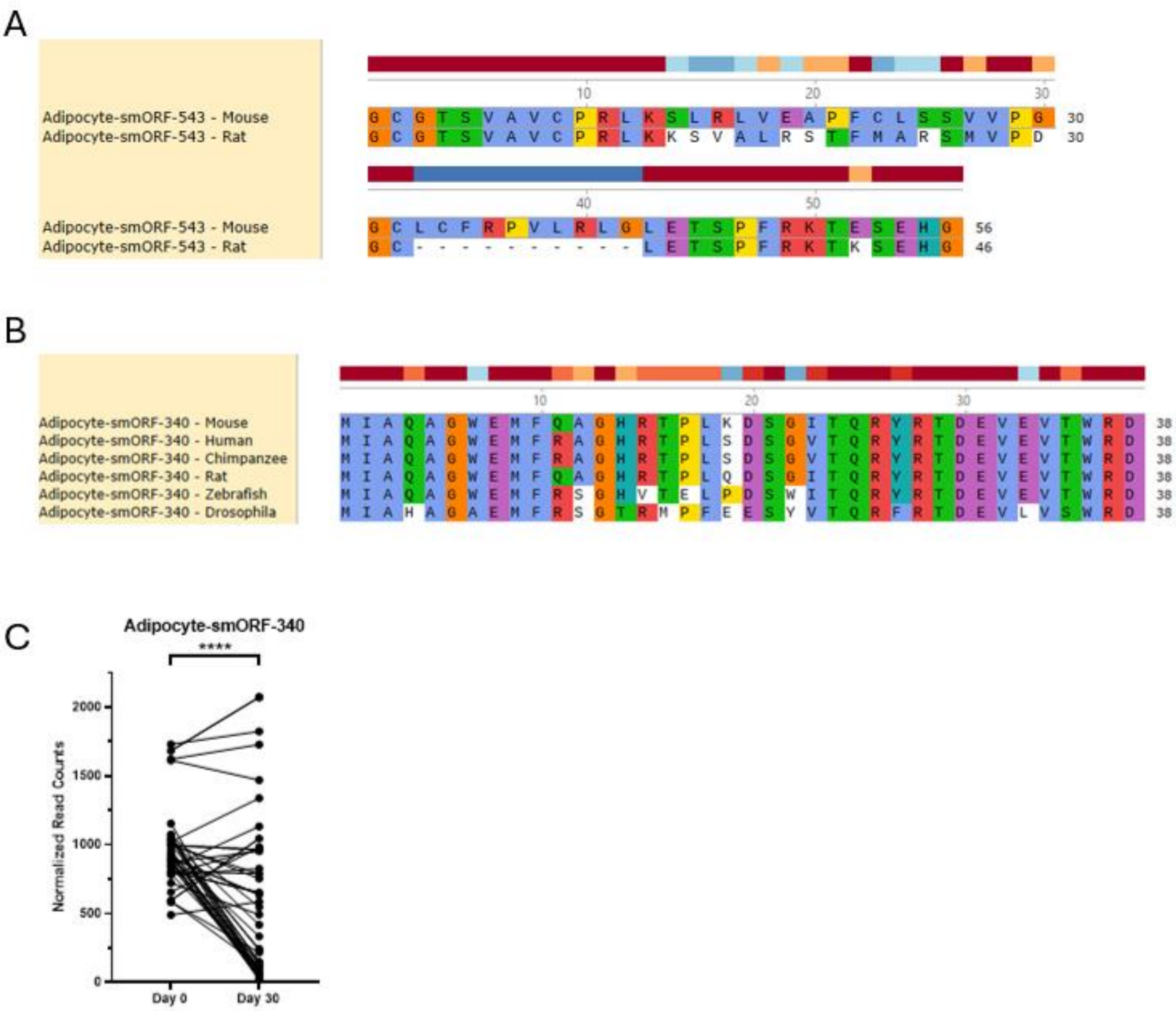


**Figure S5** Conserved and non-conserved smORFs that affect 3T3-L1 cell proliferation. **A.** Amino acid level conservation of Fat-SEP-543 as predicted by tBlastn. **B.** Amino acid level conservation of Fat-SEP-340 as predicted by tBLASTn **C.** Normalized read counts of Fat-SEP-1183 sgRNA in the CRISPR/Cas9 screen of differentiated 3T3-L1 cells. **** = P < 0.0001 by Wald’s Test calculated from MAGeCK-MLE.

Figure S6

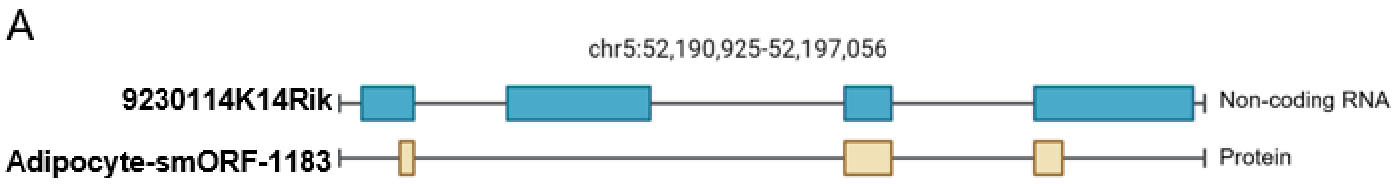


B MPCSGPLCIAMTIPGTHYVNQRSACHCLSSAGIK
VARLGSGKYSSSLVFLWIIDCTQDQKRKQLDHLQL
KNGDRNVIQVHNGLLFSYEK

**Figure S6.** Description of Fat-SEP-1183**. A.** Diagram showing predicted exons of Fat-SEP-1183. **B.** Amino acid sequence of Fat-SEP-1183.

**Supplementary Table Legends**

**Supplementary Table S1.** Excel file containing amino acid sequence, nucleotide sequence, and genome location for 3T3-L1 and SVF smORFs used in this study

**Supplementary Table S2.** Excel file containing MAGeCK-MLE analysis for CRISPR/Cas9 screen in this study. Related to Figure 3B and Figure 4A

**Supplementary Table S3**. Excel file containing tBLASTn hits for Adipocyte smORFs for Homo Sapiens, Pan Troglodytes, Rattus Norvegicus, Danio Rerio, and Drosophila Melanogaster.
